# Structure-function relationships underpin disulfide loop cleavage-dependent activation of *Legionella pneumophila* lysophosholipase A PlaA

**DOI:** 10.1101/2023.03.24.534060

**Authors:** Miriam Hiller, Maurice Diwo, Sabrina Wamp, Thomas Gutsmann, Christina Lang, Wulf Blankenfeldt, Antje Flieger

**Affiliations:** Division of Enteropathogenic Bacteria and Legionella (FG11), Robert Koch Institute, Burgstr. 37, 38855 Wernigerode, Germany; Structure and Function of Proteins, Helmholtz Centre for Infection Research, Inhoffenstrasse 7, 38124 Braunschweig, Germany; Research Center Borstel, Leibniz Lung Center, Division of Biophysics, Parkallee 10, 23845 Borstel, Germany; CSSB – Centre for Structural Systems Biology, Hamburg, Germany; Institute for Biochemistry, Biotechnology and Bioinformatics, Technische Universität Braunschweig, Braunschweig, Germany

**Author notes:** corresponding authors: Antje Flieger and Wulf Blankenfeldt and. contributed equally (Miriam Hiller, Maurice Diwo). contributed equally (Wulf Blankenfeldt, Antje Flieger).

**Keywords:** *Legionella*, phospholipase, GDSL enzyme, activation, 3D structure, disulphide loop, zinc metalloprotease

## Abstract

*Legionella pneumophila,* the causative agent of a life-threatening pneumonia, intracellularly replicates in a specialized compartment in lung macrophages, the *Legionella*-containing vacuole (LCV). Secreted proteins of the pathogen govern important steps in the intracellular life cycle including bacterial egress. Among these is the type II secreted PlaA which, together with PlaC and PlaD, belongs to the GDSL phospholipase family found in *L. pneumophila*. PlaA shows lysophospholipase A (LPLA) activity which increases after secretion and subsequent processing by the zinc metalloproteinase ProA at residue E266/L267 located within a disulfide loop. Activity of PlaA contributes to the destabilization of the LCV in the absence of the type IVB-secreted effector SdhA. We here present the 3D structure of PlaA which shows a typical α/β hydrolase fold and reveals that the uncleaved disulfide loop forms a lid structure covering the catalytic triad S30/D278/H282. This leads to reduction of both substrate access and membrane interaction before activation; however, the catalytic and membrane interaction site gets more accessible when the disulfide loop is processed. After structural modelling, a similar activation process is suggested for the GDSL hydrolase PlaC, but not for PlaD. Furthermore, the size of the PlaA substrate binding site indicated preference towards phospholipids comprising ~16 carbon fatty acid residues which was verified by lipid hydrolysis, suggesting a molecular ruler mechanism. Indeed, mutational analysis changed the substrate profile with respect to fatty acid chain length. In conclusion, our analysis revealed the structural basis for the regulated activation and substrate preference of PlaA.

## Introduction

*Legionella pneumophila* is ubiquitously found in aqueous habitats, multiplies within environmental amoebae, and is an important bacterial lung pathogen (1,2). From its natural habitat, *L. pneumophila* is transmitted via aerosols into the human lung where lung macrophages serve as the primary replication site. The infection process in mammalian cells and in amoeba shows many similarities. In both, the bacteria apply means to withstand the multifaceted host defenses and subsequently replicate within a specialized phagosome, termed the *Legionella*-containing vacuole (LCV) (3). Establishment of an intact replication vacuole and its maintenance - for the time of bacterial replication - is essential for *Legionella* propagation and requires a variety of bacterial and host factors (4,5). Several bacterial gene loci contribute to intracellular establishment and replication of *L. pneumophila* and here secretion systems, especially the type IVB (T4BSS) Dot/Icm and the type II (T2SS) Lsp systems, and their transported effectors are major determinants (6–10). It has been shown that phospholipases are among the secreted effectors and affect establishment and maintenance of the LCV. For example, patatin-like phospholipases VipD and VpdC, both exported via the type IVB Dot/Icm machinery, influence lipid composition of endosomes fusing to the LCV or vacuolar expansion by regulating lysophospholipid content, respectively (11–14). Further, phospholipases are required to exit the LCV and the host after completed replication and their involvement has been shown for a variety of pathogens including *L. pneumophila* (15–17).

Phospholipases are important virulence factors of pathogens (18) and classify into four major functional groups based on the position at which they cleave within a phospholipid, specifically phospholipases A (PLA), phospholipases B (PLB), phospholipases C (PLC), and phospholipases D (PLD). PLAs, like VipD or VpdC, hydrolyze the carboxylester bonds at sn-1 or sn-2 position and thereby release fatty acids. If only one fatty acid is targeted, a lysophospholipid is generated and may be further cleaved by a lysophospholipase A (LPLA), liberating the remaining fatty acid. At least 19 phospholipases are found in *L. pneumophila*, comprising 15 PLAs, three PLCs, and one PLD. The PLAs are divided into the patatin-like phospholipases (incl. VipD and VpdC), the PlaB-like, and the GDSL enzymes (19–21).

*L. pneumophila* possesses three GDSL enzymes PlaA, PlaC, and PlaD, on which this study is focused (Fig. 1A). The family of GDSL hydrolases is characterized by five conserved amino acid sequence blocks (19,22,23). The first block is located near the N-terminus and contains the active site serine embedded in a Gly-Asp-Ser-Leu (GDSL) motif as opposed to the Gly-X-Ser-X-Gly motif commonly found in other lipases. The catalytic triad of GDSL hydrolases is completed by aspartate and histidine located in the fifth conserved block. GDSL hydrolases have a flexible active site that allows for broad substrate specificity and enzyme activities range from protease to arylesterase to PLA/LPLA and glycerophospholipid:cholesterol acyltransferase (GCAT) activities. The latter transfers phospholipid-bound fatty acids to the acceptor molecule cholesterol (22,23).

**Figure 1:**
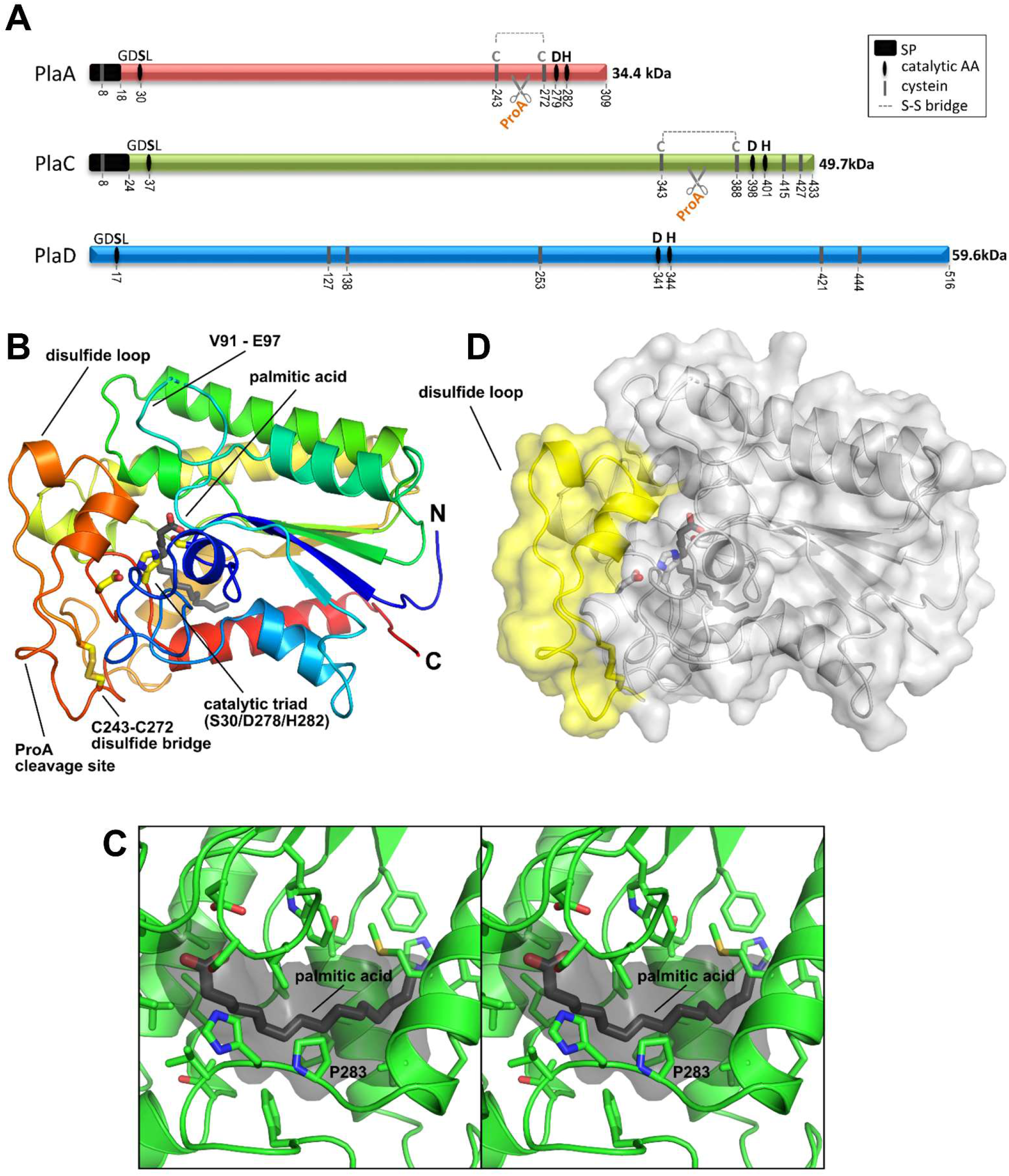
Crystal structure of *Legionella pneumophila* PlaA in complex with palmitic acid shows that the disulfide loop covers the catalytic triade. (A) Schematic overview of *L. pneumophila* PlaA, PlaC, and PlaD showing the location of the SP, the catalytic triade, the disulfide loop (only found in PlaA and PlaC) and molecular weights of unprocessed proteins Adapted from (27). ( *B*) cartoon representation of PlaA structure, colored from blue at the N-terminus to red at the C-terminus. Important residues of the active site (catalytic triad: S30, D279, H282) and the limiting residues (C243, C272) of the disulfide loop (orange) are shown as sticks (carbon: yellow, oxygen: red, nitrogen: blue, sulfur: gold), together with the product palmitic acid (carbon: black). The ProA cleavage site resides within the disulfide loop. (*C*) cross-eyed stereoplot of the PlaA substrate binding site. Palmitic acid is shown in black, the ligand binding cavity is shown in grey. Residues that contribute to the binding site are shown as sticks (carbon: green). (*D*) surface representation of PlaA. The protein is in the same orientation as in *B*. Note that the disulfide loop shown in yellow shields the active site fully. Abbreviations: SP – predicted signal peptide, AA – amino acid(s), S-S bridge – predicted disulfide bond

The three enzymes differ with regard to activities, molecular weight, and mode of secretion (Tab. 1, Fig. 1A). PlaA mainly shows LPLA and GCAT activities while PlaC reveals PLA and GCAT activities (24–26). PlaA and PlaC, which have molecular weights of 34.4 and 49.7 kDa, respectively, harbor a predicted sec signal peptide (SP) and are both secreted via the T2SS Lsp. Interestingly, the proteins observed in culture supernatants were considerably smaller after reducing SDS-PAGE, specifically ~25-28 kDa and ~36 kDa, respectively (26,27). This was however not the case for culture supernatants of a *L. pneumophila* knock out mutant in the type II-secreted zinc metalloproteinase ProA which suggested processing of PlaA and PlaC by ProA (26,27). The respective preferred cleavage sites of ProA were analyzed and are located within disulfide loop structures of both proteins. For PlaA, the preferred site was determined as E266/L267. Cleavage of the disulfide loop by ProA or disulfide loop truncation by mutagenesis is associated with an increase in enzymatic activity of PlaA and PlaC (26,27). Specifically, PlaA LPLA but not GCAT activity and PlaC PLA and GCAT activities are activated by ProA (26,27). However, the mechanism is not understood from a structure-centered view. The 59.6 kDa protein PlaD has not been characterized extensively, lacks a predicted sec secretion signal, and it is not secreted into the culture supernatant upon growth in broth medium (27).

**Table 1:** Important protein characteristics of *L. pneumophila*. Corby GDSL enzymes PlaA, PlaC, and PlaD Data are derived from this study and (26,27).NCBI Pprotein database (PlaA: WP_011947186.1; PlaC: WP_011947718.1; PlaD: WP_011947511.1).

| Protein characteristic | PlaA (lpc1811) | PlaC (lpc3121) | PlaD (lpc0558) |
| --- | --- | --- | --- |
| NCBI protein database | WP_011947186.1 | WP_011947718.1 | WP_011947511.1 |
| Size / SP [amino acids] | 309 / 18 | 433 / 24 | 519 / no SP predicted |
| MW [kDa] | 34.4 | 49.7 | 59.6 |
| MW w/o SP [kDa] | 32.8 | 46.9 | no signal peptide |
| MW w/o SP [kDa],<br>processed | 27.8 | 36.1 | not applicable |
| Preferred ProA cleavage<br>site | E266/L267 | Potential sites: **Q366/Y367**,<br>Y363/V364, Q372/Y373 | Not applicable |
| Catalytic triad | S30, D279, H282 | S37, D398, H401 | S17, D341, H344 |
| Central β-sheet | 5 parallel | 5 parallel | 5 parallel + 1 antiparallel |
| additional helices in $\alpha/\beta$-<br>hydrolase fold | No | Yes (P57-G120 and D166-P193) | No |
| Lid / disulfide loop | yes (C243-C272) | yes (C343-C388) | no lid and no disulfide loop |
| Additional C-terminal<br>domain | No | No | Yes (K400-T516) |

Relevance of PlaA and PlaC during host cell infection was shown for both enzymes. Specifically, in the absence of the T4BSS effector SdhA, which is important for LCV stabilization, PlaA can disrupt the vacuole membrane such that the bacteria get access to the cytosol, i.e. prematurely egress from the LCV and, as a consequence, host cell death pathways are triggered (15,28). Further, a *L. pneumophila* GDSL triple mutant showed reduced cell exit (17). Therefore, PlaA may be an important factor supporting bacterial LCV and host egress after intracellular replication. In line with these functions, PlaA and its activity is detected from mid-exponential growth phase on and is most prominent in late-logarithmic growth phase (29,30). Also, PlaC is more abundant during stationary phase than during logarithmic growth phase and shows a functional overlap with the aminopeptidase LapA (30,31). Although *plaC* and *lapA* individually seemed dispensable for host infections, *L. pneumophila plaC/lapA* double mutants show a severe replication defect in host amoebae (31).

Since many of the pathogen phospholipases are capable to attack cellular membranes, such enzymes may also damage the pathogen itself before export. Therefore, sophisticated activation mechanisms might be required for example to prevent self-inflicted damage to the producer or to spacially regulate activity. Such mechanisms are multifaceted and include the following: 1) oligomerization as found in *E. coli* PldA (32), 2) deoligomerization as described for *L. pneumophila* PlaB or *P. aeruginosa* PlaF (21,33,34), interaction with accessory factors, such as 3) metal ions described for *L. pneumophila* PLCs (35), 4) chemical modification, such as ubiquitination of VpdC (14), or 5) host or bacterial protein cofactors, as Rab5/Rab22 found for *L. pneumophila* VipD (11–13), and 6) proteolytic processing of proenzymes. As outlined above, the latter proteolysis-based activation is described for *L. pneumophila* PlaA and PlaC (26,27,36).

We present here the crystal structure of PlaA as basis for the clarification of structure-function relationships in disulfide loop cleavage-dependent enzyme activation. The PlaA structure is moreover compared to computational models of PlaC and PlaD to highlight similarities and differences between these enzymes. Thus, we establish a link between structure and enzymatic function of PlaA.

## Results

### PlaA displays a classical α/β-hydrolase fold containing the catalytic triad S30/D279/H282

The gene for PlaA without the respective predicted N-terminal SP (amino acids 1-18) was expressed in *E. coli* and purified via an N-terminal Strep-tag. After size exclusion chromatography (SEC), the protein was subjected to crystallization. PlaA crystallized in spacegroup P3_1_21 with one chain of the monomeric protein in the asymmetric unit. This crystal form differs from a previous report where PlaA has been crystalized in space groups P2_1_ and P2_1_2_1_2_1_ but where structure determination has remained pending (37). Initial phases were obtained from single-wavelength anomalous diffraction data collected at an X-ray energy of 7 keV (wavelength = 1.7712 Å) of a crystal obtained in the presence of ammonium iodide (Fig. S1A and B). Crystallization in the presence of 1-monopalmitoyllysophosphatidylcholine (16:0 LPC) and avoiding polyethylene glycol (PEG) in the precipitant led to crystals of the same spacegroup (Fig. S1A) and no significant structural differences were observed between the crystal structures determined in this study. With the exception of a few residues at the N- and C-termini as well as parts of a highly flexible surface loop (residues V91 – E97), the final structures show all of the PlaA protein (Fig. 1B). The N-terminal Strep-tag was too flexible to produce traceable electron density. The highest resolution obtained here is 1.45 Å for PlaA complexed with the 16:0 LPC hydrolysis product palmitate (Tab. 2). The structures substantiate that PlaA is a canonical GDSL hydrolase with a typical α/β-hydrolase fold (Fig. 1B, Fig. S1B). The catalytic triad was previously determined as S30, D279, H282 by means of mutational analysis (27) and is confirmed in the 3D structure. The central β-sheet consists of five parallel β-strands and is surrounded by a set of α-helices on both of its sides (Fig. 1B, Fig. S1C). The N-terminus and the C-terminus of PlaA (shown in blue and red in Fig. 1B) are located in close proximity to each other.

**Table 2:**
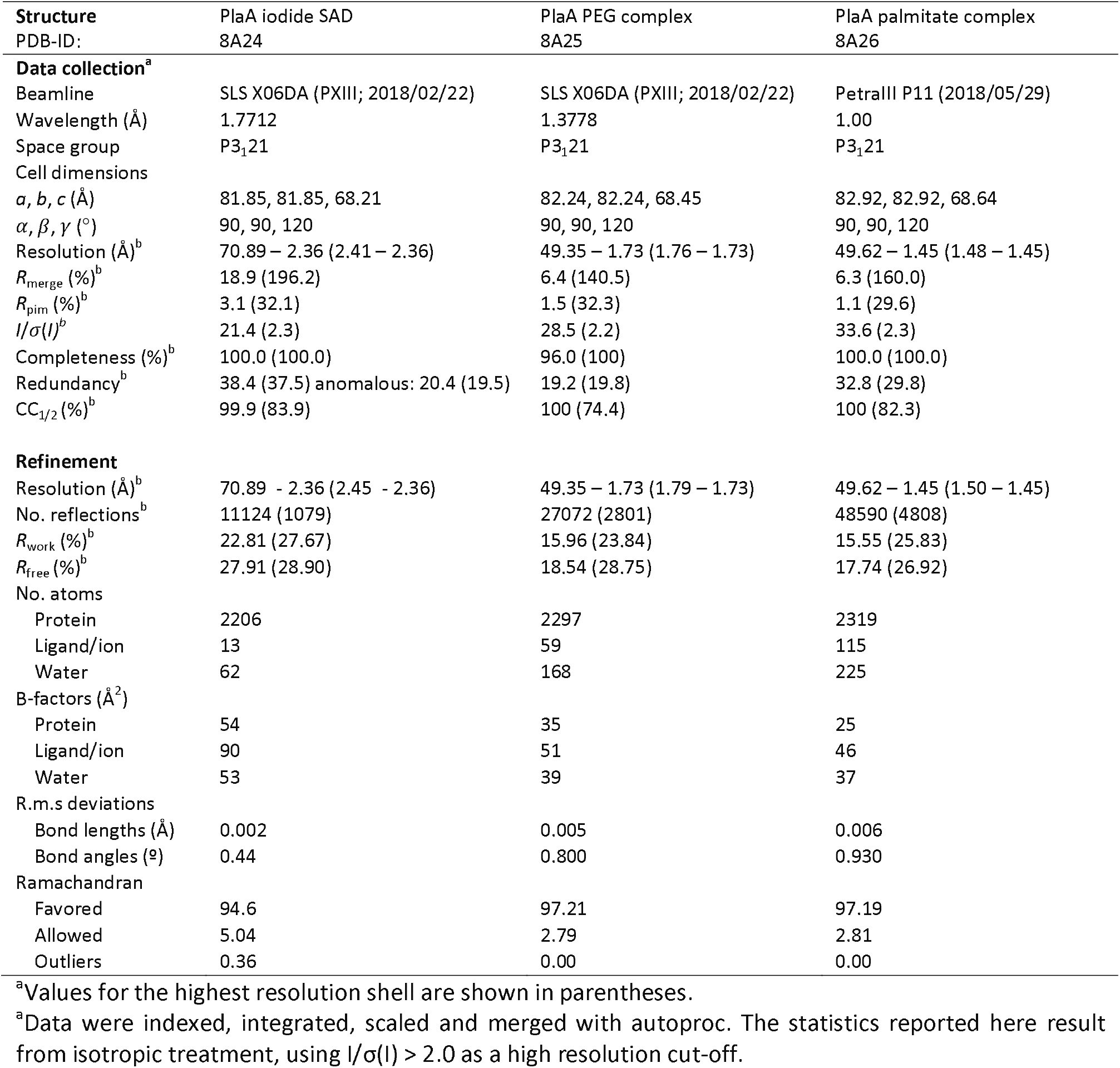
X-ray data collection and refinement statistics.

### Detection of a hydrophobic channel that matches with linear fatty acids of 16 carbon atoms in close proximity to the catalytic triad

Interestingly, crystallization was initially only possible in the presence of PEG. Here, we observed additional electron density in a hydrophobic channel in close proximity to the catalytic triad (Fig. S1D), despite the fact that this channel is not accessible from the solvent. The channel has a length of approx. 20 Å, suggesting that the unexplained electron density derives from a molecule that mimics the fatty acid moiety of the substrate such as short unbranched PEG components of the precipitant used to crystallize the protein. We hence interpreted the additional electron density as a fragment of PEG. Subsequently, we aimed for the structure of a complex of PlaA with its substrate 16:0 LPC by co-crystallization in a PEG-free precipitant. This yielded crystals of the same space group (Fig. S1A) that diffracted to 1.45 Å resolution. Here, we observed additional electron density that occupies the entire hydrophobic channel (Fig. S1E). This density was interpreted as the product of PlaA-dependent hydrolysis of 16:0 LPC, palmitic acid (Fig. 1C). Interestingly, no crystals were obtained in the absence of substrate (16:0 LPC) or product analogues (palmitate, PEG), which may point towards higher flexibility of apo-PlaA. This flexibility may include partial unfolding for hydrophobic substrate binding.

### A disulfide loop functions as a lid that covers the catalytic triad and reduces membrane interaction before enzyme activation

Regarding potential structural elements hindering substrate binding in the non-activated state of PlaA, a disulfide loop (from C243 to C272; shown in orange in Fig. 1B and in yellow in Fig. 1D, respectively) covers the entrance to the substrate binding pocket and the catalytic triad of PlaA. It is therefore conceivable that ProA-mediated processing of the disulfide loop facilitates access of substrates to the binding pocket and to the active site, explaining the increased LPLA activity upon exposure to ProA. The previously determined preferred proteolytic maturation site E266/L267 at the C-terminal end of the lid (27) is solvent accessible in the crystal structure (Fig. 1B, Fig. S1C). Parts of the lid display high B-factors, indicating that they are relatively mobile. This flexibility may also explain why we found the hydrolysis product palmitic acid bound to the active site (Fig. 1C and Fig. S1E) despite having crystallized uncleaved PlaA. We propose that proteolytic maturation by ProA increases flexibility further and enables parts of the lid to leave the position observed in the crystal structure. This would open up the entry to the active site and is in line with proteolytic activation mechanisms of other lipases (38).

Among others, DALI (39) searches identify the cholin esterase ChoE from *P. aeruginosa* (PDB code 6uqv) (40), the phospholipase PlpA from *Vibrio vulnificus* (PDB code 6jl2) (41), or the esterase domain of autotransporter EstA from *P. aeruginosa* (PDB code 3kvn) (42) as highly structurally similar to PlaA (Fig. 2A-D). Interestingly, while these proteins also possess disulfide loops in similar positions, these disulfide loops are shorter in sequence and they do not shield the active center as much as the disulfide loop of PlaA in the unprocessed state observed here (Fig. 2A-D). As a consequence, the ligand binding sites of these enzymes are solvent accessible, suggesting that they do not require proteolytic activation.

**Figure 2:**
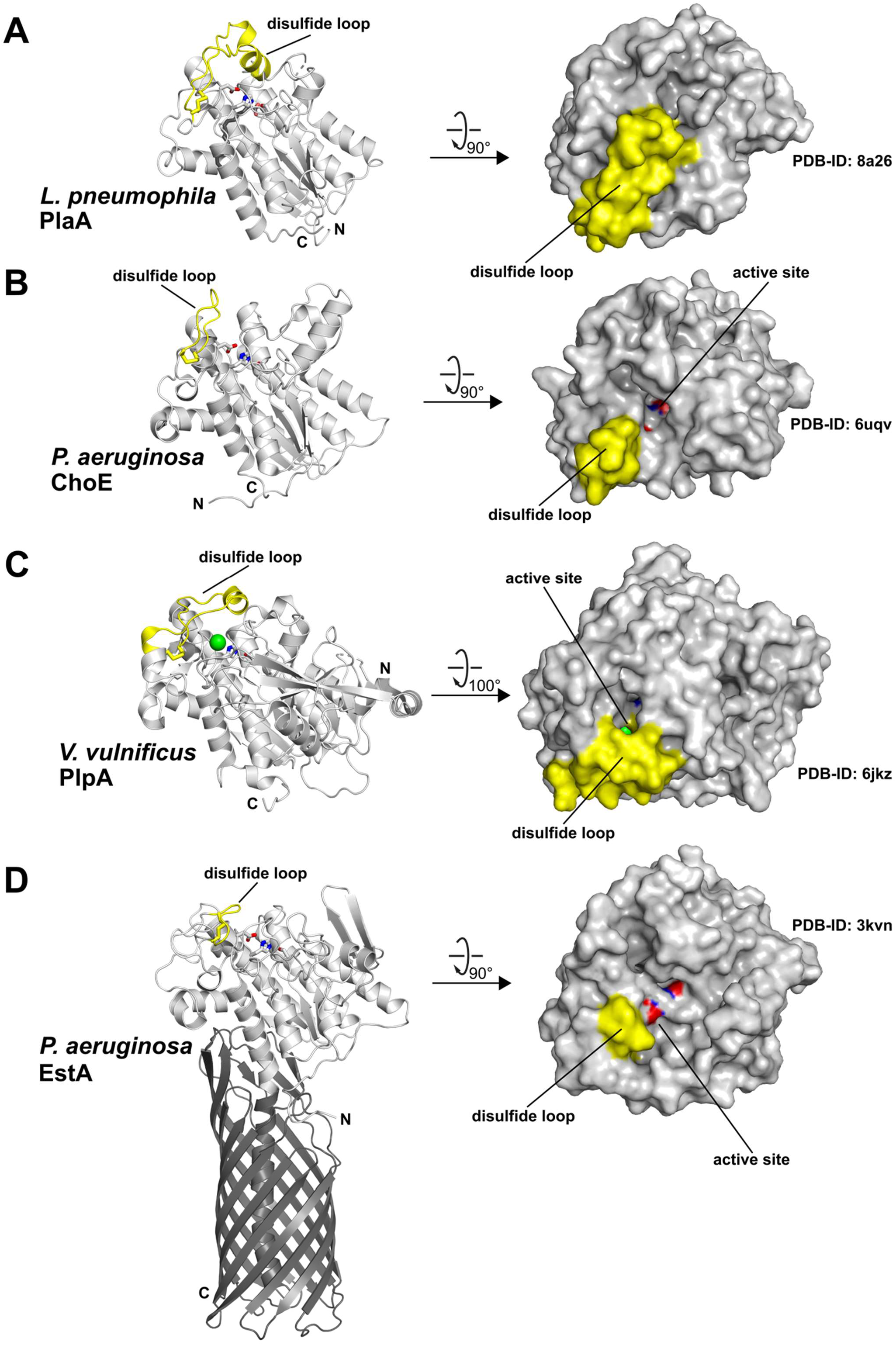
Crystal structures of selected proteins related to *L. pneumophila* PlaA according to analysis with DALI (39). Ribbon representations with residues of the catalytic triad in the active site are shown as sticks on the left; top views of the molecular surface are shown on the right. Residues of the disulfide loops are plotted in yellow. Oxygen and nitrogen atoms of the catalytic triad are shown in red and blue, respectively. (*A*) Crystal structure of *L. pneumophila* PlaA in complex with palmitic acid. (*B*) *P. aeruginosa* ChoE is an acetylcholinesterase (PDB entry 1uqv) (40). (*C*) *V. vulnificus* PlpA is a type II - secreted phospholipase A2 that utilizes a chloride ion (green sphere) instead of aspartate in its catalytic triad (PDB entry 6jkz) (41). (*D*) EstA is an autotransported esterase displayed on the outer membrane of *P. aeruginosa*. The autotransporter domain is shown in darker colors (PDB entry 3kvn) (42). Note that ChoE, PlpA and EstA have significantly shorter disulfide loops than PlaA, which makes their active sites solvent-accessible.

We further analyzed whether in addition to the previously shown increase in activity (27) also membrane interaction of PlaA is influenced by disulfide loop processing and tested the catalytic inactive variant PlaA S30N versus the respective mutant with truncated disulfide loop PlaA S30N Δaa248-267, in a Förster-Resonance Energy Transfer (FRET) liposome-insertion assay. When protein-lipid interaction exceeds peripheral binding and proteins at least insert into the lipid headgroup region, this leads to an increase in the distance between the fluorophores and thus to reduced energy transfer. Consequently, the ratio of donor and acceptor fluorescence intensities is a measure of protein insertion into membranes. This effect is very prominent for antimicrobial peptides, like LL-37, and leads to a strong increase in the ratio (43). For the disulfide loop mutant PlaA S30N ΔΔaa248-267 a concentration-dependent increase of the ratio was as well detected and was higher compared to PlaA S30N with intact disulfide loop (Fig. 3). However, the interaction of the disulfide loop mutant was less pronounced as noted for LL-37 (43). This indicates that the PlaA disulfide loop mutant only inserts into the interface of the membrane. In conclusion, our results show that the disulfide lid reduces lipid interaction; or in other words, lipid interaction is facilitated by disulfide loop cleavage which might also impact enzyme activity.

**Figure 3:**
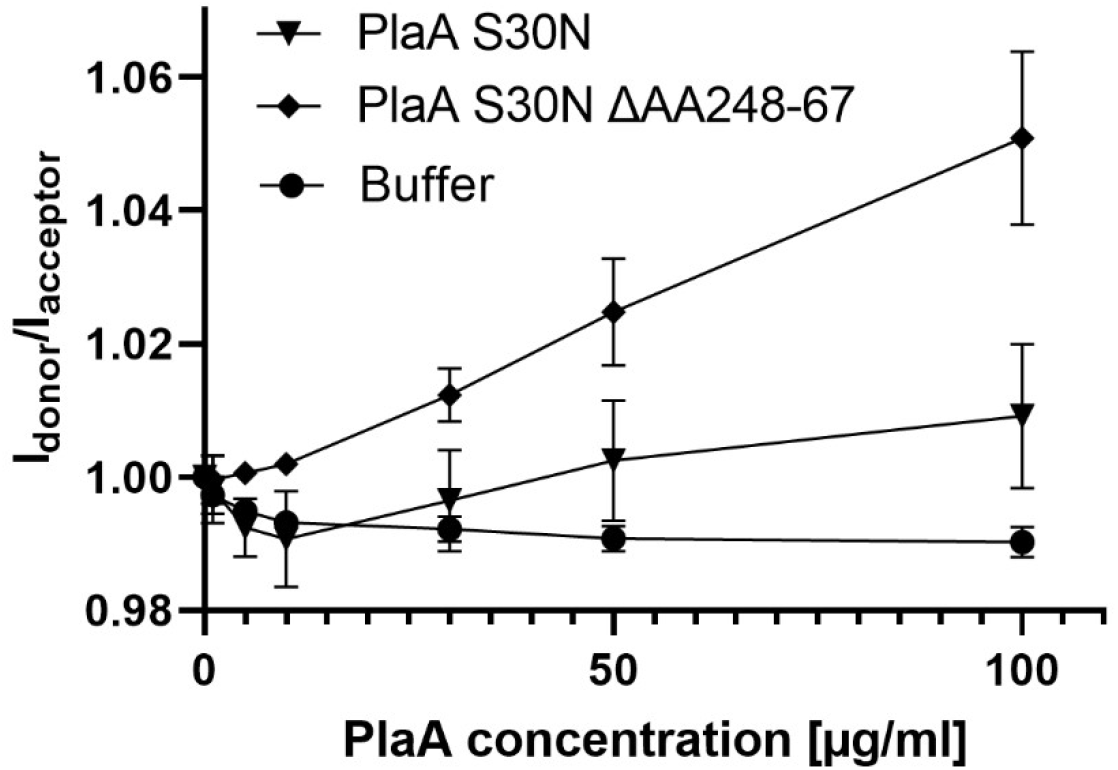
Deletion of the disulfide loop in PlaA increases membrane interaction. Protein-membrane interaction of PlaA S30N and disulfide loop deletion mutant PlaA S30N ΔAA248-267 was monitored by FRET spectroscopy. An increase of the intensity ratio I_donor_/I_acceptor_ corresponds to a reduced FRET efficiency and indicates insertion of proteins into the ePC:ecPE (1:1) liposomes. Experiments were performed in 40 mM Tris, 150 mM NaCl, pH 7.5, at 37 °C. Donor and acceptor intensities of the liposomes were both adjusted to 300,000 cps. PlaA variant concentrations of 1, 5, 10, 30, 50, and 100 µg/ml were titrated into the liposome solution and the respective values were determined 5 min after injection when a steady state was reached. Experiments were repeated four times and the data are shown as means and standard deviation (n=4).

### Substrate selectivity of PlaA concerning fatty acid chain length is governed by a molecular ruler mechanism

As mentioned above, we observed unexplained elongated electron density in the ligand binding site of the first high-resolution structure of PlaA and attributed this to a PEG fragment deriving from the precipitant solution (Fig. S1D). However, this molecule did not fill the binding site completely, and further structural analysis suggested that PlaA can accommodate phospholipid substrates with linear fatty acid chains of at least 16 carbon atoms length, which was then confirmed by the crystal structure of PlaA in complex with palmitate (Fig. 1C, Fig. S1E). To further investigate the substrate spectrum of PlaA, we tested whether LPLA activity of PlaA correlated with the length of the fatty acid chains. To this end, PlaA was incubated with lysophosphatidylcholine (LPC) with diverse fatty acid residues. These varied in length from eight to 22 carbon atoms. PlaA in its unprocessed form exhibited predominantly activity towards 16:0 LPC (Fig. 4A, left side). Additionally, activity was compared to the disulfide loop-deleted PlaA ΔAA248-67, which corresponds to ProA-processed PlaA (27). The LPLA activity of PlaA ΔAA248-67 was distinctly higher than that of untruncated PlaA and was mainly directed towards 16:0 and 14:0 LPC, but 8:0 LPC, 12:0 LPC, and 18:0 LPC were also hydrolyzed to a lower extent (Fig. 4A, right side). As hypothesized, no activity towards 20:0 LPC and 22:0 LPC was observed for both PlaA proteins. PlaA variants with mutation of the catalytic serine Ser30 (PlaA S30N and PlaA S30N ΔAA248-67) were used as controls and did not exhibit activity towards any of the tested lipid substrates (Fig. 4A, left and right side, respectively). These findings suggest that the substrate selectivity of PlaA is governed by a molecular ruler mechanism that measures the length of fatty acids by accommodating them to the ligand binding site (44). To corroborate this, we mutated residue P283 pointing into the ligand binding site to leucine (Fig. 1C and Fig. S2), hypothesizing that this larger residue would diminish the hydrophobic lipid binding site and hence change the substrate preference of PlaA to lysophospholipids with shorter fatty acid chains (Fig. 1C and Fig. S3). Indeed, although showing an overall reduced activity, PlaA P283L preferred 12:0 LPC over 14:0 and 16:0 LPC (Fig. 4B), highlighting the importance of the size of this ligand binding site. In contrast, PlaA T220L where another residue in the binding site was mutated to leucine which however did not point as much into the ligand binding site like P283L and accordingly did not show a shift in the substrate preference (Fig. 4B and Fig. S2). In summary, our data suggest that substrate preference with regard to fatty acid chain length is regulated by a ruler like mechanism.

**Figure 4:**
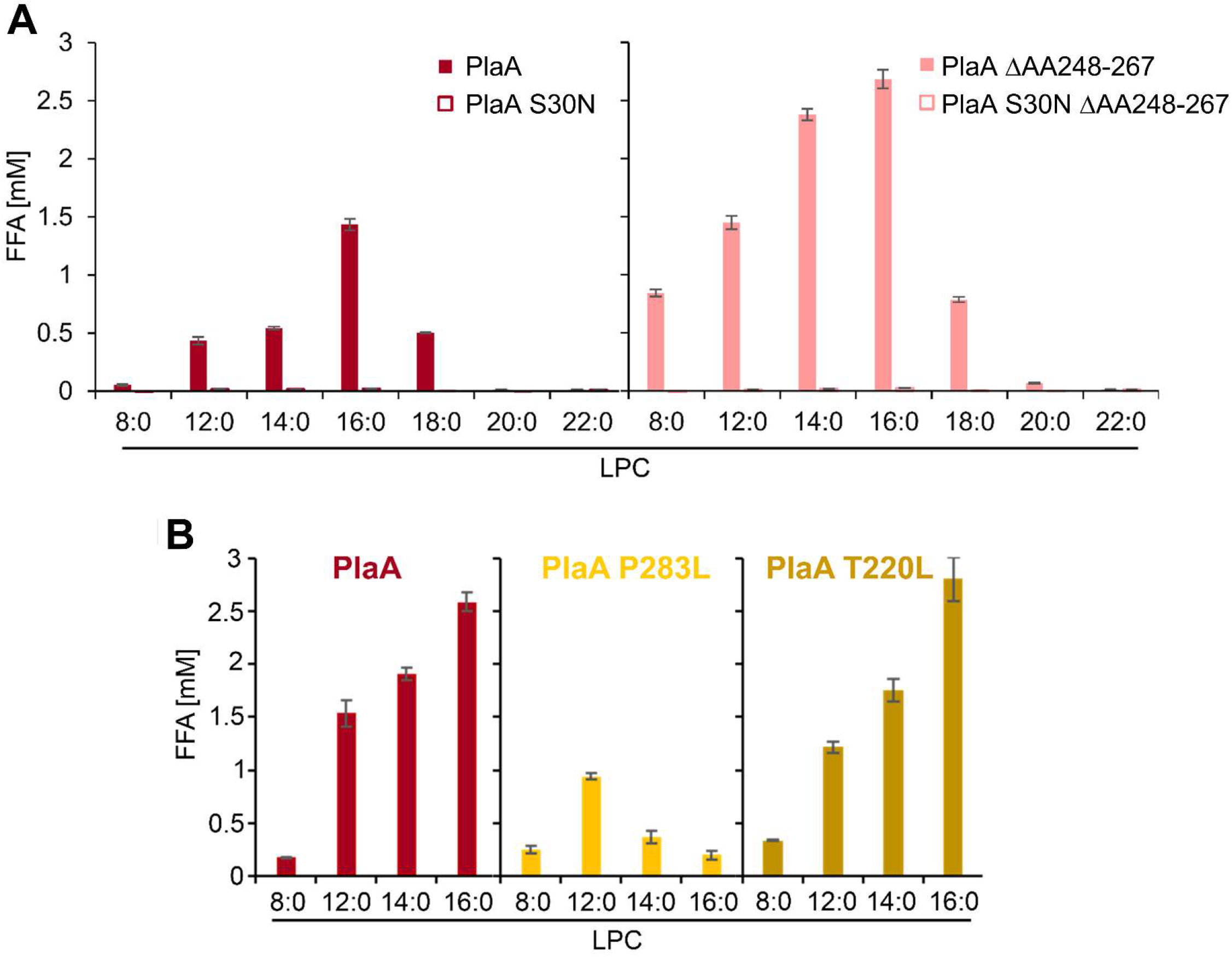
PlaA prefers 16:0 lysophospholipids for hydrolysis and a P283L mutation in the substrate binding pocket changes substrate preference to shorter fatty acids. (*A, B*)-LPLA activities of PlaA and PlaA variants were determined via quantification of free fatty acids (FFA) after incubation for 111h with the indicated lipids at 3711°C. (A) 200ng recombinant protein of PlaA and its catalytic inactive mutant PlaA S30N (left side) or a disulfide loop truncation mutant PlaA ΔΔAA248-67 and its catalytic inactive mutant PlaA S30N ΔΔAA248-67 (right side) were used for the assay. (B) Cell lysates of strains expressing PlaA and PlaA mutants in the substrate binding site, such as PlaA P283L or PlaA T220L, were used and diluted 1:2. Location of the mutated residue P283L is shown in Fig. 1C and mutated residues P283L and T220L in Fig. S2. Data are shown as means and standard deviation (n=3) and are representative for at least two additional experiments.

### During growth of *L. pneumophila*, PlaA is first present in its unprocessed form until ProA is secreted

PlaA is a type II-secreted protein found in culture supernatants of *L. pneumophila* (24,25,27). We were interested if and when ProA-processing of PlaA occurs in *L. pneumophila* grown in liquid cultures. PlaA was detected as ~33 kDa full-length protein (without signal peptide, protein sizes see Tab. 1) during lag and logarithmic growth phases with a maximum quantity at 2 to 3 h of growth (Fig. 5A and B). During mid-exponential growth, from ~4 h of incubation, processed PlaA with a molecular weight of ~25 kDa was detected for the first time and from 5h remained the only PlaA version until late logarithmic / stationary phase (Fig. 5A and 5B). These observations correlated with an accumulation of ProA in the culture supernatants over time (Fig. 5C). In conclusion, our experiments indicate that PlaA, although found in its full-length form in the culture supernatant at early growth phases, may not reach its full activity before its processing at mid to late logarithmic growth phase. Full activity of PlaA may therefore correlate to a later phase in bacterial growth and be associated with LCV and host cell exit of the bacteria, as indicated previously (15).

**Figure 5:**
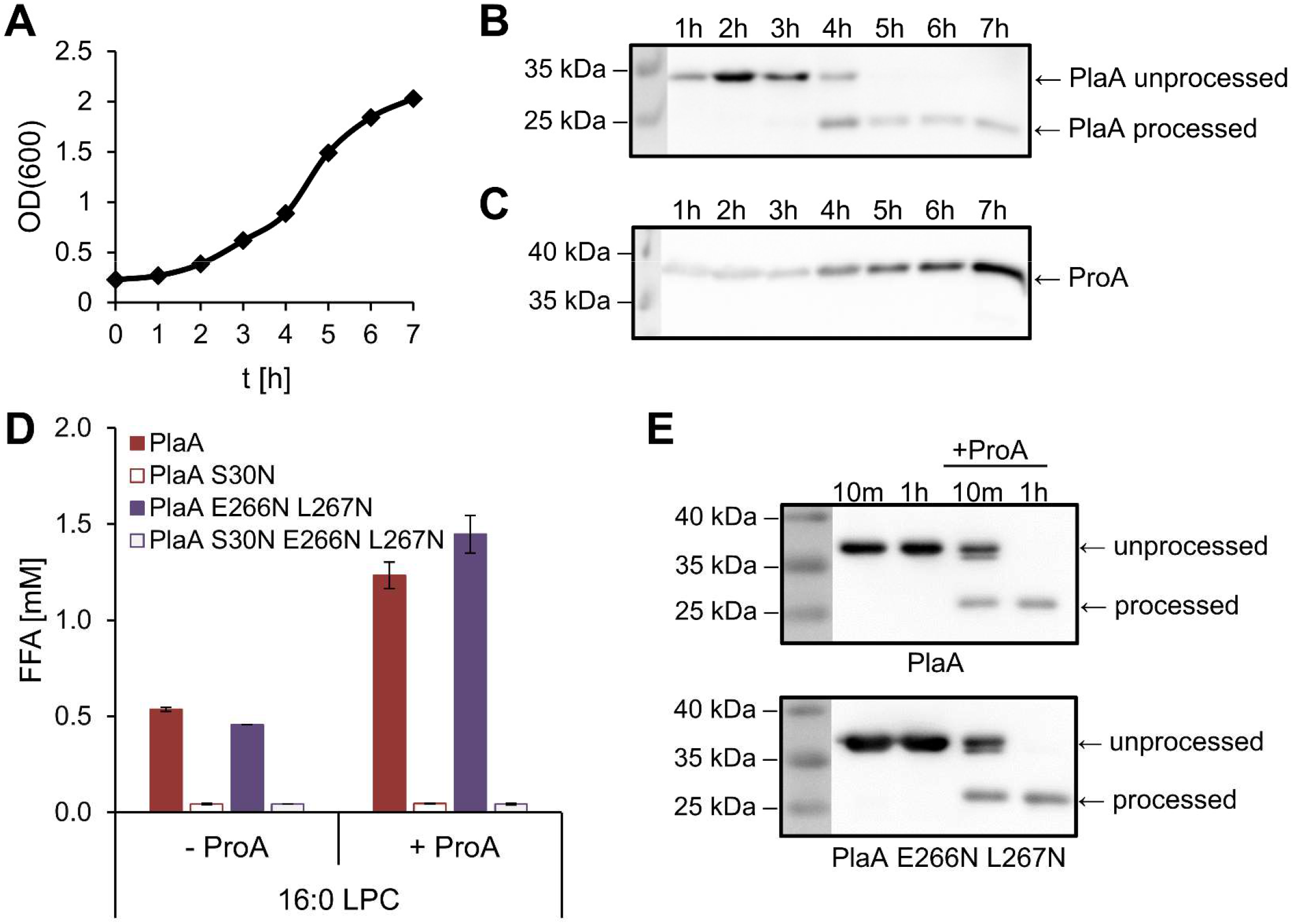
Processing of PlaA occurs from mid-logarithmic growth and PlaA activation by ProA is not inhibited by mutation of E266/L267 highlighting ProA’s broad cleavage site acceptance. *(A)* The growth of *L. pneumophila* wild type in broth was monitored over a period of 7 h and samples of culture supernatants were analyzed for the presence, processing status of PlaA *(B)*, and presence of ProA *(C)* by Western blotting with PlaA- and ProA-directed antibodies, respectively. Molecular weight standards are shown on the left of the Western blots. The data are representative for one additional experiment. (*D*) LPLA activities of recombinant PlaA (25 ng), a PlaA variant mutated in the preferred cleavage site E266N L267N and catalytic inactive S30N mutants of both were determined via quantification of free fatty acids (FFA) after incubation for 311h with 16:0 LPC at 3711°C. Reactions were performed without or with addition of 3.5 mU ProA as indicated. (*E*) Western Blots using PlaA-directed antibodies after incubation of PlaA and a mutant in the preferred cleavage site E266N L267N without and with addition of ProA for 10 min and 1 h. ProA-derived processing occurred for both types of PlaA. (*D*) Data are shown as mean and standard deviation (n=3) and (D and E) are representative for three additional experiments.

### PlaA activation by ProA- is not inhibited by mutation of the preferred cleavage site E266/L267 highlighting broader cleavage site acceptance by ProA

As outlined above, an important feature of PlaA is a disulfide loop lid structure. After its processing access of lipid substrates to the catalytic active site and protein-lipid interaction is facilitated and thus leads to an increase of LPLA activity. We previously reported that this disulfide loop is cleaved by ProA at the preferred cleavage site E266/L267 in PlaA. Also, truncation of the disulfide loop in PlaA ΔAA248-67 yields more active PlaA and similar activity characteristics as ProA-processed PlaA (27). Subsequently, we analyzed whether mutation of the main cleavage site E266/L267 may hinder PlaA activation. We found that the mutation of E266 and L267 into N266 and N267, respectively, is not sufficient to prevent ProA-dependent increase in LPLA activity of PlaA and its proteolytic processing. Specifically, similar activities were detected for PlaA and the E266N L267N mutant and similar processing to the smaller version of PlaA by ProA (Fig. 5D and 5E). The respective catalytic inactive S30N controls did not show activity (Fig. 5D). This indicates that ProA may attack other exposed residues surrounding the major cleavage site E266/L267 in the disulfide loop and lead to PlaA processing and activation. This was already implied by peptide mapping of ProA-digested PlaA which identified additional but less preferred cleavage sites, such as K264/P265, P265/E266, and L267/T268, directly upstream and downstream of E266/L267 (27). These observations further highlight the previously described broad cleavage site acceptance by ProA in other substrates (45).

### Comparison of PlaA to structural models of PlaC and PlaD highlights the presence of disulfide loops shielding the catalytic side in PlaC but not PlaD

To obtain additional insight into the structure of the three *L. pneumophila* GSDL phospholipases, we generated structural models of PlaC and PlaD using the template-free routine of AlphaFold (46). The structure of PlaA was predicted as well to evaluate the method, and no significant differences were found with respect to the X-ray structures described above (Fig. 6A, Fig. S3A-C). Notably, this also includes the more flexible parts of PlaA (residues V91 – E97 and the disulfide loop), whose flexibility is correctly indicated by a lower confidence score. Importantly, the predicted conformation of the disulfide loop matches the crystal structure very well (Fig. S3A-C).

**Figure 6:**
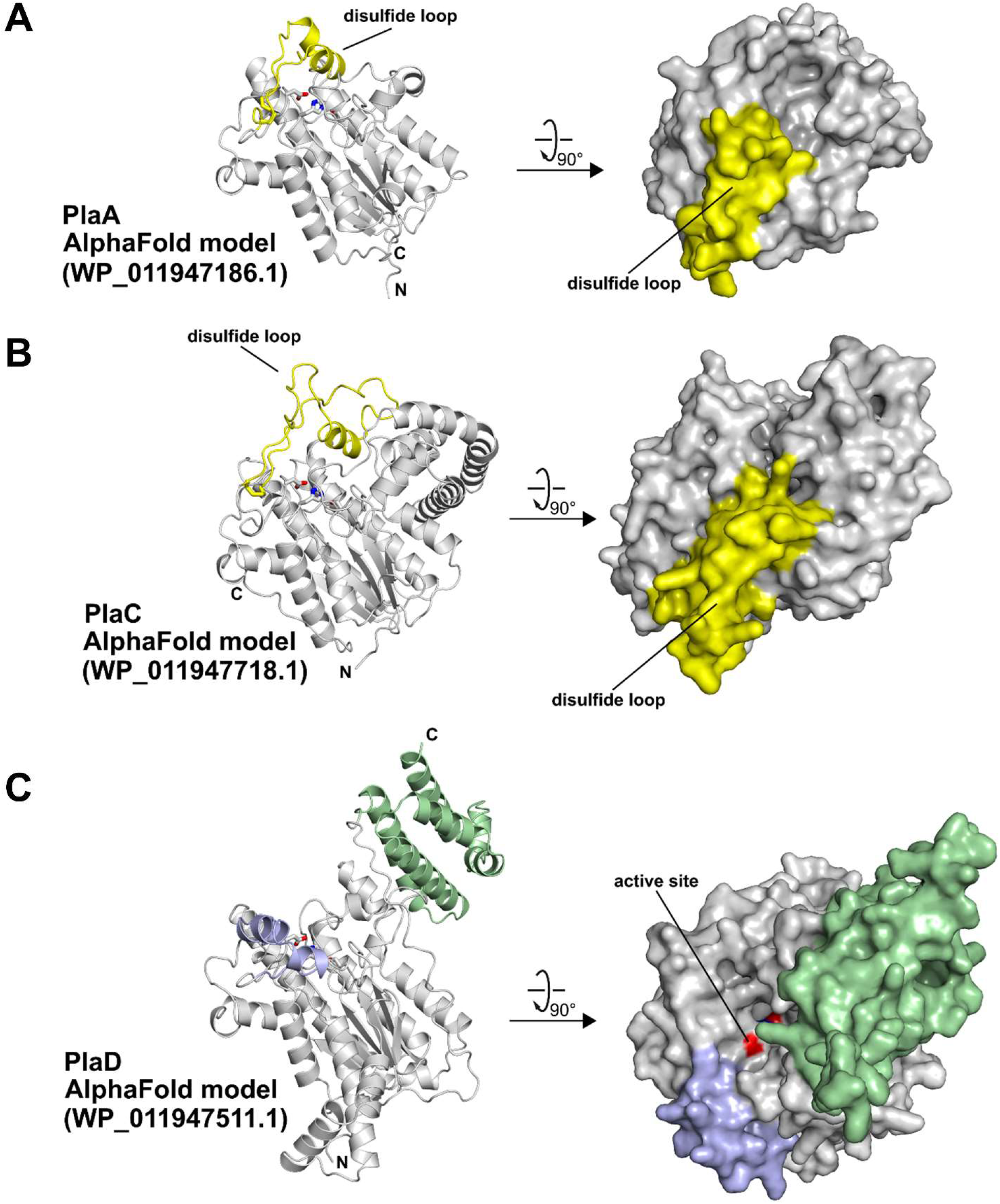
AlphaFold (46) predicted structures of the three *L. pneumophila* GDSL phospholipases PlaA, PlaC, and PlaD. Representations, orientations and color schemes are similar to Fig. 2. (*A*) AlphaFold model of PlaA from *L. pneumophila* strain Corby (NCBI Protein database entry WP_011947186.1). Note the correct prediction of the position of the disulfide loop with respect to the experimental structure (compare to Fig. 2A and Fig. S2A-C), leading to a shielded active site. (*B*) AlphaFold model of PlaC from *L. pneumophila* strain Corby (NCBI Protein database entry WP_011947718.1). The predicted position of the disulfide loop leads to a similar shielding as in PlaA. (*C*) AlphaFold model of PlaD from *L. pneumophila* strain Corby (NCBI Protein database entry WP_011947511.1). PlaD does not contain a disulfide loop; residues structurally corresponding to the disulfide loop of PlaA are shown in light blue. The predicted additional C-terminal domain of unknown function of PlaD is shown in light green. Note that the active site of PlaD is predicted to be solvent-exposed, however, the C-terminal domain may position itself to block access to the active center.

Compared to PlaA, PlaC although similar in structure is predicted to contain additional helical elements in P57 – G120 and D166 – P193 which correspond to its larger size (Fig. 6B). Of these, the second region replaces the flexible region V91 – D95 of PlaA. Similar to PlaA, PlaC has a lid domain that is delimited by a disulfide bridge (C343-C388) which accommodates the proteolytic maturation site (26). In PlaC, this lid includes an α-helix (362-370) that shields the active site from the solvent. However, this α-helix is slightly shifted away from the active site, resulting in a larger entrance funnel for PlaC possibly explaining differences in the substrate specificity of both enzymes. While PlaA preferentially hydrolyzes lysophospholipids, PlaC also accepts larger substrates such as diacyl phospholipids (26). The length of PlaC’s hydrophobic substrate channel is identical to PlaA’s, indicating that in both cases linear fatty acids with ~16 carbon atoms are the preferred substrates.

The structure of PlaD includes the α/β hydrolase domain (N-terminal domain) and is predicted to have an additional α-helical domain of unknown function at its C-terminus (P403 – T516) (Fig. 6C). The relative orientation towards the α/β-hydrolase domain is unclear at present since both domains are connected by a long flexible linker (P371 - L402). Comparison with PlaA reveals additional helical elements (e.g. P60 – P72). Importantly, PlaD does not contain a disulfide loop, and the respective region of the protein (Y311 – A334, shown in light blue in Fig. 6C) is predicted to adopt an open conformation that leads to an unshielded substrate binding site. In addition to differences in protein secretion, these features may explain further possible differences in the activation mechanism of this protein compared to PlaA and PlaC. The length of the hydrophobic substrate channel of PlaD, on the other hand, is identical to that of PlaA, again suggesting preference for linear fatty acids with ~16 carbon atoms.

## Discussion

Here we show that PlaA, which is one of three proteins of the GDSL phospholipase family found in *L. pneumophila* (19), adopts an α/β hydrolase fold typically found in similar enzymes of other bacteria, such as the cholin esterase ChoE from *Pseudomonas aeruginosa* (PDB code 6uqv), the phospholipase PlpA from *Vibrio vulnificus* (PDB code 6jl2) or the esterase domain of autotransporter EstA from *P. aeruginosa* (PDB code 3kvn) to name just a few examples (Fig. 2). However, PlaA also shows specific features which relate to its activation mechanism and substrate specificity. Structural analysis reveals that a disulfide loop lid structure reduces substrate access to the catalytic site in ProA-unprocessed PlaA. The active site gets more accessible when the disulfide loop is opened up by proteolysis. After structural modelling using AlphaFold, a similar activation mechanism is suggested for PlaC and cleavage of the disulfide loop and PlaC activation by ProA was previously shown (26) but is not suggested for PlaD, where the catalytic site seemed freely accessible as further observed in ChoE, PlpA, and EstA (Fig. 2).

Lid structures are very common in lipases and are important for substrate specificity and control of enzymatic activity (47). It can be assumed that processing of the PlaA disulfide loop by ProA might completely remove the lid or at least increase its flexibility. This might affect the conformation of the active site or the potential substrate-binding domain or both. It is conceivable that the access of lipid substrates to the active site channel of PlaA is facilitated by the structural alterations induced by ProA-dependent processing, which might explain the increased LPLA activity of processed PlaA. Proteolytic processing is a common mechanism for activation of enzymes that have been produced as inactive zymogens (38). Among others, the induction of enzymatic activity after processing within a disulfide loop has been shown for some, such as *Pseudomonas* exotoxin A and Shiga toxin from enterohemorrhagic *Escherichia coli* (48–51). Although unprocessed PlaA is not completely inactive, it can be assumed that the rationale of ProA-dependent regulation of PlaA is to increase the enzymatic activity only after PlaA is secreted to the exterior and at the required time point. Since phospholipase activities are potentially harmful to the bacterium itself, we propose that the here presented activation mechanism is also used to avoid self-inflicted lysis from a lytic enzyme before secretion.

*L. pneumophila* is an intracellularly replicating lung pathogen which induces major lung damage leading to a severe pneumonia and the secreted bacterial phospholipases may contribute to the clinical picture (2,19). Eukaryotic membranes and the surfactant phospholipid monolayer in the lung contain major amounts of phospholipids with esterified fatty acids of a length of ~16 carbon atoms (52). We showed that the size of an intramolecular channel in PlaA, PlaC, and PlaD nicely fits to harbor such lipids and also the reaction product palmitic acid (Fig. 1C) which mechanistically underlines host-adapted substrate specificity of these enzymes.

Interestingly, it was shown that PlaA is responsible for the destabilization of the replication vacuole in the absence of the T4BSS secreted effector SdhA, which localizes to the LCV (15). The LCV protective mechanism of SdhA has recently been shown as binding and blocking the function of OCRL (OculoCerebroRenal syndrome of Lowe), an inositol 5-phosphatase pivotal for controlling endosomal dynamics and participating in vacuole disruption (28). It is important for intracellular pathogens like *Legionella* to maintain the integrity of the replication vacuole for the period of bacterial propagation to avoid detection by the immune system of the host (55). However, for the exit of the pathogen and the initiation of a new infection cycle, the vacuole needs to be disrupted (16). Assuming that the functions of SdhA and PlaA are opposed and must be balanced for maintenance of the LCV integrity for bacterial replication (55), it may be that a change of abundance or activity of either factor could result in exit of *L. pneumophila*. The here addressed increase of PlaA activity and membrane interaction by proteolytic processing could disturb the balance between SdhA and PlaA in an infection and might thus lead to destabilization of the LCV triggering the exit process.

Our analyses of culture supernatants at multiple time points during *L. pneumophila* growth in liquid medium revealed that PlaA is initially secreted as full-length protein. Processing was first detected during mid-exponential growth when ProA quantities increased. We speculate that ProA-processing of PlaA does not occur before a critical density of intravacuolar bacteria is reached which produces sufficient ProA quantities. This might happen in similarity to the regulatory process of quorum sensing (56). In parallel, proteolytic activation of PlaC might be additionally triggered which might allow cooperation of PlaC, majorly revealing PLA activity, and PlaA, majorly possessing LPLA activity. Thus, destabilization of the LCV for host exit can be linked to coordinated activation of several enzyme activities at high cell density and completed bacterial replication.

In summary, these findings provide insight into the relationship between structure and activity of PlaA and for the other GSDL hydrolases in *L. pneumophila*. We found that these enzymes possess structural features that may lead to a specific activity and activation profile tailored to the needs of the bacterium.

## Experimental procedures

### Bacterial strains and growth conditions

All experiments and sequence or structure predictions were performed with *L. pneumophila* sg1 strain Corby including recombinant gene expression in *E. coli* BL21. The strains used and generated in the study are listed in Tab. 3. *L. pneumophila* strains were grown on buffered charcoal yeast extract (BCYE) agar and in buffered yeast extract (BYE) broth and *E. coli* strains were grown on Luria Bertani (LB) agar and in LB broth as described previously (36,57,58). If appropriate, 100 µg/ml ampicillin or 50 µg/ml kanamycin were added to the *E. coli* cultures.

**Table 3:**
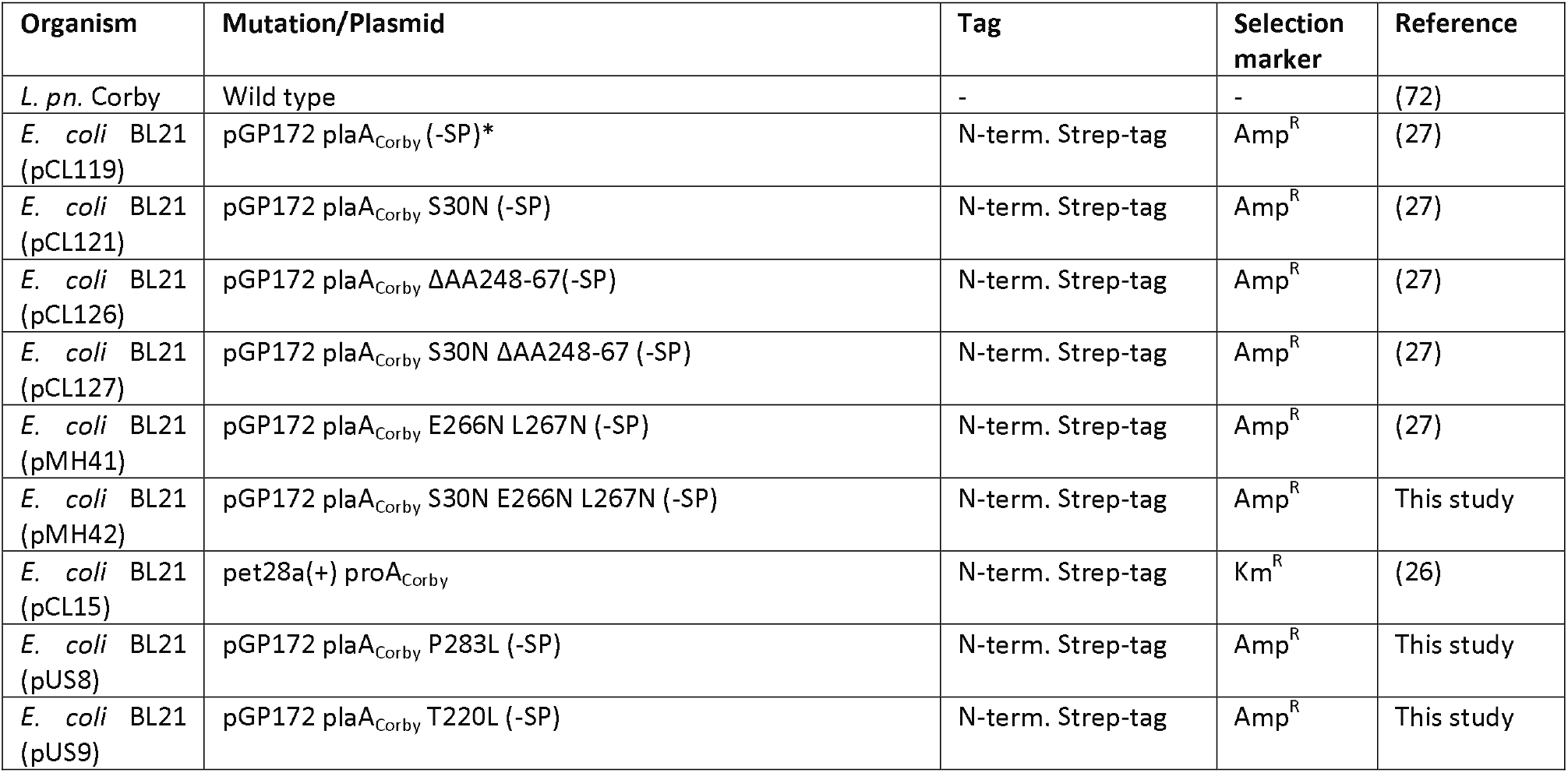
Overview of strains used in this study. *-(SP): constructs cloned without sequence coding for the predicted signal peptide (aa 1-18), strain was used for PlaA expression and purification for crystallization, X-ray diffraction, and structure determination.

### Construction of plasmids

The vector pMH42 was generated from pMH41 by means of the QuikChange Site-Directed Mutagenesis Kit (Stratagene) using the primers PlaA_S30N_fw (5‘-GTATTTGGTGATAATTTGTCGGATAACGG-3‘) and PlaA S30N_rv (5‘-CCGTTATCCGACAAATTATCACCAAATAC-3‘) for introduction of the S30N point mutation in *plaA* (Tab. 3). The vectors pUS8 and pUS9 were generated from pCL119 using the primer pairs plaA_P283L_fw/rv (5’-GATTTGGTTCATCTGACAGCG-*3’/%’-CGCTGTCAGATGAACCAAATC-3’) and plaA_P220L_fw/rv (5’-GACTTGGGTAGTCATTTTGACC-3’/5 GAC’TACCC-AAGTCAAAAAACAAC-3’), respectively (Tab. 3). Primers were obtained from Eurofins MWG Operon and Integrated DNA Technologies. All restriction enzymes were obtained from New England Biolabs. Plasmid DNA was introduced into *E. coli* by heat shock transformation.

### Expression and purification of recombinant PlaA or ProA for enzyme activity tests and analysis of crystal structure

*E. coli* BL21 strains harboring plasmids for recombinant expression of *plaA* variants without SP and N-terminally fused Strep-tag (Tab. 3) were grown in LB broth supplemented with ampicillin at 37 °C and 250 rpm to an OD_600_ = 0.8, induced with 0.1 mM IPTG, and transferred to 18 °C and 250 rpm for 16 – 17 h. The bacterial pellet was collected and resuspended in 100 mM Tris-HCl, pH 8.0, containing 100 mM NaCl. Homogenization was performed with Emulsi Flex C3 (Avestin) applying 25,000 to 30,000 psi. The soluble fraction was collected by centrifugation at 13,000 g and 4 °C for 60 min. Strep-PlaA and variants were purified using Strep-Tactin Superflow high capacity resin (IBA) and further purified by SEC (Superdex S200 10/30 Increase) performed with SEC buffer (20 mM Tris pH 8, 150 mM NaCl, 1 mM DTT) according to the manufacturer’s instructions. Recombinant periplasmic ProA was isolated by osmotic shock from *E. coli* BL21 (pCL15) and subsequent anion exchange chromatography (26).

### Crystallization, X-ray diffraction, and structure determination

Crystals were obtained by sitting drop vapor diffusion at 20 °C. Initially, 200 nl PlaA solution at 8 mg/mL in SEC buffer were mixed with 200 nl of 14.7% (w/v) PEG3350, 1% (w/v) PEG 1K, 0.5% (w/v) PEG200, 8.9% (v/v) tacsimate pH 5, 200 mM NH_4_I, 1 mM CaCl_2,_ and 1 mM MgCl_2_. For cryoprotection, crystals were briefly washed in a solution of 10% (v/v) (2R,3R)-(-)-2,3-butandiole, 16% (w/v) PEG3350, 6% (v/v) Tacsimate pH 5, and 920 mM NH_4_I. To exploit the f’/f’’ difference arising from the L I absorption edge of iodine (5.1881 keV, 2.3898 Å), two datasets at an X-ray energy of 7.000 keV (λ = 1.771196 Å) and with detector distances of 140 mm and 135 mm were collected at beamline X06DA (PXIII) of the Swiss Light Source (SLS; Villigen, Switzerland) (59). These data were indexed, scaled and merged with autoproc (60) and then used for initial phasing by single anomalous dispersion (SAD) analysis through hkl2map and the SHELX-package (61) The resulting poly-alanine model served as a template for phenix.autobuild (62). Manual editing was done in Coot (63) from the CCP4 software suite (64). Structure refinement was done with phenix.refine and pdb-redo (65,66). 13 iodide anions were placed in the final model (Fig. S1B). High resolution data of a crystal obtained under similar conditions were collected at the same beamline, using an X-ray wavelength of 1.3778 Å. Diffraction data of a crystal grown from 7 mg/mL PlaA solution in SEC buffer supplemented with 500 µM 1-monopalmitoyllysophosphatidylcholine (16:0 LPC) with a precipitant consisting of 2.2 M (NH_4_)_2_SO_4_ and 0.2 M NH_4_-acetate were collected on beamline P11 of the PETRAIII synchrotron (DESY Hamburg, Germany) (67). The cryoprotectant consisted of precipitant solution supplemented with 10% (v/v) (2R,3R)-(-)-2,3-butandiole in this case. Refinement proceeded via Fourier synthesis followed by employing the same methods as in the case for the SAD structure (specific data see Tab. 2).

### Prediction of 3D structures of PlaA, PlaC and PlaD

The 3D structures of PlaA (gene lpc1811), PlaC (gene lpc3121) und PlaD (gene lpc0558) of *L. pneumophila* Corby were predicted using the template-less routine of AlphaFold (46) through a local installation of ColabFold (68). The required protein sequences were obtained from the NCBI protein database (PlaA: WP_011947186.1; PlaC: WP_011947718.1; PlaD: WP_011947511.1).

### Preparation of cell lysates

Pellets of 1 ml overnight cultures of strains expressing PlaA and PlaA variants were lysed with 0.5 mg/ml lysozyme, 1% Triton X-100 in 1 ml 40 mM Tris-HCl buffer and supplemented with 6 mM NaN_3_.

### *In vitro* analysis of lipolytic activity and proteolytic processing of recombinant PlaA

Lipolytic activities of recombinant PlaA variants were investigated as described previously (25,36,69). The lipid substrates 1-monooctanoyllysophosphatidylcholine (8:0 LPC), 1-monolauroyllysophosphatidylcholine (12:0 LPC), monotetradecanoyllysophosphatidylcholine (14:0 LPC), 1-monopalmitoyllysophosphatidylcholine (16:0 LPC), 1-monostearoyllysophosphatidylcholine (18:0 LPC), 1-monoarachidoyllysophosphatidylcholine (20:0 LPC), and 1-monobehenoyllysophosphatidylcholine (22:0 LPC) purchased from Avanti Polar Lipids (Alabaster, AL, USA) were applied for the assays. Briefly, purified recombinant proteins or cell lysates expressing recombinant protein were incubated with 6.7 mM lipid suspensions as micelles (in 40 mM Tris-HCl, 1% Triton X-100) with or without addition of 3.5 mU ProA at 37 °C for the indicated time followed by quantitative detection of free fatty acids (FFA) by means of the NEFA-C kit (WAKO Chemicals, Neuss, Germany) according to the instructions of the manufacturer.

### FRET liposome-interaction assay

Insertion of PlaA proteins into liposome membranes was determined in 40 mM Tris, 150 mM NaCl, pH 7.5, at 37 °C by FRET spectroscopy applied as a probe dilution assay (70). L-α-phosphatidylcholine (ePC, 95%, chicken egg) and L-α-phosphatidylethanolamine (ecPE, *E. coli*) were purchased from Avanti (Avanti Polar Lipids Inc., Alabaster, AL, USA) and prepared at stock concentrations of 10 mg/mL in CHCl_3_. Fluorescence conjugated lipids *N*-(7-nitrobenz-2-oxa-1,3-diazol-4-yl)-1,2-dihexadecanoyl-*sn*-glycero-3-phosphoethanolamine (NBD-PE) and Lissamine^TM^ Rhodamine B 1,2-dihexadecanoyl-*sn*-glycero-3-phosphoethanolamine (Rho-DHPE), were purchased from Life Technologies (Carlsbad, CA, USA). Proteins were added to PC:PE (1:1) liposomes, which were labelled with 1% of the donor NBD-PE and 1% of the acceptor rhodamine-PE. The fluorescence was measured by means of a spectrometer (Fluorolog 3, Horiba Jobin Yvon GmbH, Bensheim, Germany). NBD excitation wavelength was 470 nm. Protein insertion was monitored as the increase of the ratio of the donor fluorescence intensity I_donor_ at 531 nm to that of the acceptor intensity I_acceptor_ at 593 nm in a concentration dependent manner. This ratio depends on the Förster efficiency; therefore, a ratio > 1 highlights that the mean distance of donor and acceptor dyes increased. Values were determined 5 min after addition of 1, 5, 10, 30, 50, and 100 µg/ml protein. Buffer (see above) was used to dilute the proteins and control.

### Analysis of processing status of PlaA from *L. pneumophila*

Overnight cultures of *L. pneumophila* sg1 strain Corby grown in buffered yeast extract (BYE) broth were washed once with warm BYE broth to remove secreted proteins, adjusted to OD_600_ = 0.3 in 37 °C BYE broth and incubated for 8 h at 37 °C and 250 rpm. OD_600_ was measured hourly, aliquots of the cultures were adjusted to OD_600_ = 0.1 and supernatants were concentrated 100-fold by trichloroacetic acid precipitation. For processing analysis of PlaA by ProA, 250ng PlaA or PlaA E266N L267N were incubated with 0.5 mM ProA for 10-60 min at 37°C. These samples were analyzed in Western blots with primary polyclonal rabbit antibodies against PlaA and ProA (generated by BioGenes from recombinant proteins) (27).

## Figure preparation

Figures were prepared with Microsoft Power Point 2010 and Inkscape 0.92. All images, symbols and fonts within the figures are acquired from Microsoft Power Point 2010. Images of crystal structures were generated with PyMOL (71). Western Blots were labelled and processed with Adobe Photoshop CS6. Graphs included in the Figures are visualized by Microsoft Excel 2010.

## Data availability statement

All data generated or analyzed during this study are included in this published article or in the supplementary information. Structures are available at PDBe with the accession codes 8A24 (PlaA iodide SAD), 8A25 (PlaA complex with PEG fragment), and 8A26 (PlaA complex with palmitate).

## Supporting information

This article contains supporting information.

## Acknowledgements

We thank Susanne Karste, Ute Strutz, and Christian Galisch (Robert Koch-Institut) and Sabrina Groth (Research Center Borstel) for excellent technical assistance. AF, TG, and WB, received funding from the German Research Association (DFG) (FL359/6-1/2; priority program SPP2225 Exit Strategies of Intracellular Pathogens FL359/10-1 and GU568/8-1; 281361126/GRK2223). We acknowledge the staff at beamlines PXIII/X06DA at SLS (Paul Scherrer Institute, Villigen, Switzerland) and P11 (PETRAIII, DESY Hamburg, Germany) for providing access to their facilities.

## Authors contributions

conceptualized the study: M.H., M.D., C.L., W.B., A.F.

supervised the study: W.B., A.F.

conceived and designed the experiments: M.H., M.D., S.W., C.L., W.B., A.F., T.G.

acquired data: M.H., M.D., S.W., C.L.

conducted the experiments: M.H., M.D., S.W., C.L.

analyzed the data: M.H., M.D., S.W., C.L., W.B., A.F., T.G.

interpreted the results: M.H., M.D., S.W., C.L., W.B., A.F., T.G.

drafted the manuscript: M.H., M.D., W.B., A.F.

revised the manuscript: M.H., M.D., S.W., C.L., W.B., A.F., T.G.

approved the final version: M.H., M.D., S.W., C.L., W.B., A.F., T.G.

## Data sharing

All data generated or analyzed during this study are available in this article and the appendices.

## Abbreviations

CTD: C-terminal domain
ePC: L-α-phosphatidylcholine (chicken egg)
ecPE: L-α-phosphatidylethanolamine (*E. coli*)
FFA: free fatty acids
FRET: Förster-Resonance-Energy-Transfer
GCAT: glycerophospholipid:cholesterol acyltransferase
LCV: *Legionella*-containing vacuole
LPLA: lysophospholipase A
NBD-PE: *N*-(7-nitrobenz-2-oxa-1,3-diazol-4-yl)-1,2-dihexadecanoyl-*sn*-glycero-3-phosphoethanolamine
MW: molecular weigth
PEG: polyethylene glycol
PLA/PLB/PLC/PLD: phospholipase A/B/C/D
Rho-DHPE: Lissamine^TM^ Rhodamine B 1,2-dihexadecanoyl-*sn*-glycero-3-phosphoethanolamine
SAD: single anomlalous dispersion
SEC: size exclusion chromatography
SP: sigal peptnide
T2SS/T4SS: type II/IV secretion system
8:0 LPC: 1-monooctanoyllysophosphatidylcholine
12:0 LPC: 1-monolauroyllysophosphatidylcholine
14:0 LPC: 1-monotetradecanoyllysophosphatidylcholine
16:0 LPC: 1-monopalmitoyllysophosphatidylcholine
18:0 LPC: 1-monostearoyllysophosphatidylcholine
20:0 LPC: 1-monoarachidoyllysophosphatidylcholine
22:0 LPC: 1-monobehenoyllysophosphatidylcholine

## References

1. Fields, B. S. (1996) The molecular ecology of legionellae. Trends Microbiol. 4, 286–290

2. Mondino, S., Schmidt, S., Rolando, M., Escoll, P., Gomez-Valero, L., and Buchrieser, C. (2020) Legionnaires’ Disease: State of the Art Knowledge of Pathogenesis Mechanisms of Legionella.

3. Horwitz, M. A. (1983) The Legionnaires’ disease bacterium (Legionella pneumophila) inhibits phagosome-lysosome fusion in human monocytes. The Journal of Experimental Medicine 158, 2108–2126

4. Finsel, I., and Hilbi, H. (2015) Formation of a pathogen vacuole according to Legionella pneumophila: how to kill one bird with many stones. Cell. Microbiol. 17, 935–950

5. Isberg, R. R., O’connor, T. J., and Heidtman, M. (2009) The Legionella pneumophila replication vacuole: making a cosy niche inside host cells. Nature Reviews Microbiology 7, 13–24

6. Segal, G., Purcell, M., and Shuman, H. A. (1998) Host cell killing and bacterial conjugation require overlapping sets of genes within a 22-kb region of the Legionella pneumophila genome. Proc. Natl. Acad. Sci. U. S. A. 95, 1669–1674

7. Cianciotto, N. P. (2009) Many substrates and functions of type II secretion: lessons learned from Legionella pneumophila. Future Microbiol. 4, 797–805

8. White, R. C., and Cianciotto, N. P. (2019) Assessing the impact, genomics and evolution of type II secretion across a large, medically important genus: the Legionella type II secretion paradigm. Microbial genomics 5

9. Burstein, D., Amaro, F., Zusman, T., Lifshitz, Z., Cohen, O., Gilbert, J. A., Pupko, T., Shuman, H. A., and Segal, G. (2016) Genomic analysis of 38 Legionella species identifies large and diverse effector repertoires. Nat. Genet. 48, 167

10. Gomez-Valero, L., Rusniok, C., Carson, D., Mondino, S., Pérez-Cobas, A. E., Rolando, M., Pasricha, S., Reuter, S., Demirtas, J., and Crumbach, J. (2019) More than 18,000 effectors in the Legionella genus genome provide multiple, independent combinations for replication in human cells. Proceedings of the National Academy of Sciences 116, 2265–2273

11. Gaspar, A. H., and Machner, M. P. (2014) VipD is a Rab5-activated phospholipase A1 that protects Legionella pneumophila from endosomal fusion. Proc. Natl. Acad. Sci. U. S. A. 111, 4560–4565

12. VanRheenen, S. M., Luo, Z. Q., O’Connor, T., and Isberg, R. R. (2006) Members of a Legionella pneumophila family of proteins with ExoU (phospholipase A) active sites are translocated to target cells. Infect. Immun. 74, 3597–3606

13. Shohdy, N., Efe, J. A., Emr, S. D., and Shuman, H. A. (2005) Pathogen effector protein screening in yeast identifies Legionella factors that interfere with membrane trafficking. Proc. Natl. Acad. Sci. U. S. A. 102, 4866–4871

14. Li, X., Anderson, D. E., Chang, Y. Y., Jarnik, M., and Machner, M. P. (2022) VpdC is a ubiquitin-activated phospholipase effector that regulates Legionella vacuole expansion during infection. Proc. Natl. Acad. Sci. U. S. A. 119, e2209149119

15. Creasey, E. A., and Isberg, R. R. (2012) The protein SdhA maintains the integrity of the Legionella-containing vacuole. Proc. Natl. Acad. Sci. U. S. A. 109, 3481–3486

16. Flieger, A., Frischknecht, F., Hacker, G., Hornef, M. W., and Pradel, G. (2018) Pathways of host cell exit by intracellular pathogens. Microb Cell 5, 525–544

17. Striednig, B., Lanner, U., Niggli, S., Katic, A., Vormittag, S., Brulisauer, S., Hochstrasser, R., Kaech, A., Welin, A., Flieger, A., Ziegler, U., Schmidt, A., Hilbi, H., and Personnic, N. (2021) Quorum sensing governs a transmissive Legionella subpopulation at the pathogen vacuole periphery. EMBO Rep 22, e52972

18. Flores-Diaz, M., Monturiol-Gross, L., Naylor, C., Alape-Giron, A., and Flieger, A. (2016) Bacterial Sphingomyelinases and Phospholipases as Virulence Factors. Microbiol. Mol. Biol. Rev. 80, 597–628

19. Hiller, M., Lang, C., Michel, W., and Flieger, A. (2018) Secreted phospholipases of the lung pathogen Legionella pneumophila. Int. J. Med. Microbiol. 308, 168–175

20. Lang, C., and Flieger, A. (2011) Characterisation of Legionella pneumophila phospholipases and their impact on host cells. Eur. J. Cell Biol. 90, 903–912

21. Diwo, M., Michel, W., Aurass, P., Kuhle-Keindorf, K., Pippel, J., Krausze, J., Wamp, S., Lang, C., Blankenfeldt, W., and Flieger, A. (2021) NAD(H)-mediated tetramerization controls the activity of Legionella pneumophila phospholipase PlaB. Proc. Natl. Acad. Sci. U. S. A. 118

22. Akoh, C. C., Lee, G.-C., Liaw, Y.-C., Huang, T.-H., and Shaw, J.-F. (2004) GDSL family of serine esterases/lipases. Prog. Lipid Res. 43, 534–552

23. Upton, C., and Buckley, J. T. (1995) A new family of lipolytic enzymes? Trends Biochem. Sci. 20, 178–179

24. Flieger, A., Gong, S., Faigle, M., Stevanovic, S., Cianciotto, N. P., and Neumeister, B. (2001) Novel lysophospholipase A secreted by Legionella pneumophila. J. Bacteriol. 183, 2121–2124

25. Flieger, A., Neumeister, B., and Cianciotto, N. P. (2002) Characterization of the gene encoding the major secreted lysophospholipase A of Legionella pneumophila and its role in detoxification of lysophosphatidylcholine. Infect. Immun. 70, 6094–6106

26. Lang, C., Rastew, E., Hermes, B., Siegbrecht, E., Ahrends, R., Banerji, S., and Flieger, A. (2012) Zinc metalloproteinase ProA directly activates Legionella pneumophila PlaC glycerophospholipid:cholesterol acyltransferase. J. Biol. Chem. 287, 23464–23478

27. Lang, C., Hiller, M., and Flieger, A. (2017) Disulfide loop cleavage of Legionella pneumophila PlaA boosts lysophospholipase A activity. Sci. Rep. 7, 16313

28. Choi, W. Y., Kim, S., Aurass, P., Huo, W., Creasey, E. A., Edwards, M., Lowe, M., and Isberg, R. R. (2021) SdhA blocks disruption of the Legionella-containing vacuole by hijacking the OCRL phosphatase. Cell Rep 37, 109894

29. Flieger, A., Gong, S., Faigle, M., Northoff, H., and Neumeister, B. (2001) In vitro secretion kinetics of proteins from Legionella pneumophila in comparison to proteins from non-pneumophila species. Microbiology 147, 3127–3134

30. Aurass, P., Gerlach, T., Becher, D., Voigt, B., Karste, S., Bernhardt, J., Riedel, K., Hecker, M., and Flieger, A. (2016) Life Stage-specific Proteomes of Legionella pneumophila Reveal a Highly Differential Abundance of Virulence-associated Dot/Icm effectors. Mol. Cell. Proteomics 15, 177–200

31. White, R. C., Gunderson, F. F., Tyson, J. Y., Richardson, K. H., Portlock, T. J., Garnett, J. A., and Cianciotto, N. P. (2018) Type II Secretion-Dependent Aminopeptidase LapA and Acyltransferase PlaC Are Redundant for Nutrient Acquisition during Legionella pneumophila Intracellular Infection of Amoebas. MBio 9

32. Dekker, N. (2000) Outer-membrane phospholipase A: known structure, unknown biological function: MicroReview. Mol. Microbiol. 35, 711–717

33. Kuhle, K., Krausze, J., Curth, U., Rossle, M., Heuner, K., Lang, C., and Flieger, A. (2014) Oligomerization inhibits Legionella pneumophila PlaB phospholipase A activity. J. Biol. Chem. 289, 18657–18666

34. Bleffert, F., Granzin, J., Caliskan, M., Schott-Verdugo, S. N., Siebers, M., Thiele, B., Rahme, L., Felgner, S., Dormann, P., Gohlke, H., Batra-Safferling, R., Jaeger, K. E., and Kovacic, F. (2022) Structural, mechanistic, and physiological insights into phospholipase A-mediated membrane phospholipid degradation in Pseudomonas aeruginosa. Elife 11

35. Aurass, P., Schlegel, M., Metwally, O., Harding, C. R., Schroeder, G. N., Frankel, G., and Flieger, A. (2013) The Legionella pneumophila Dot/Icm-secreted effector PlcC/CegC1 together with PlcA and PlcB promotes virulence and belongs to a novel zinc metallophospholipase C family present in bacteria and fungi. J. Biol. Chem. 288, 11080–11092

36. Banerji, S., Bewersdorff, M., Hermes, B., Cianciotto, N. P., and Flieger, A. (2005) Characterization of the major secreted zinc metalloprotease-dependent glycerophospholipid:cholesterol acyltransferase, PlaC, of Legionella pneumophila. Infect. Immun. 73, 2899–2909

37. Qu, X., Song, X., Zhang, N., Ma, J., and Ge, H. (2020) The phospholipase A effector PlaA from Legionella pneumophila: expression, purification and crystallization. Acta Crystallogr F Struct Biol Commun 76, 138–144

38. Vanaman, T. C., and Bradshaw, R. A. (1999) Proteases in cellular regulation minireview series. J. Biol. Chem. 274, 20047–20047

39. Holm, L. (2020) DALI and the persistence of protein shape. Protein Sci. 29, 128–140

40. Pham, V. D., To, T. A., Gagne-Thivierge, C., Couture, M., Lague, P., Yao, D., Picard, M. E., Lortie, L. A., Attere, S. A., Zhu, X., Levesque, R. C., Charette, S. J., and Shi, R. (2020) Structural insights into the putative bacterial acetylcholinesterase ChoE and its substrate inhibition mechanism. J. Biol. Chem. 295, 8708–8724

41. Wan, Y., Liu, C., and Ma, Q. (2019) Structural analysis of a Vibrio phospholipase reveals an unusual Ser-His-chloride catalytic triad. J. Biol. Chem. 294, 11391–11401

42. van den Berg, B. (2010) Crystal Structure of a Full-Length Autotransporter. J. Mol. Biol. 396, 627–633

43. de Miguel Catalina, A., Forbrig, E., Kozuch, J., Nehls, C., Paulowski, L., Gutsmann, T., Hildebrandt, P., and Mroginski, M. A. (2019) The C-Terminal VPRTES Tail of LL-37 Influences the Mode of Attachment to a Lipid Bilayer and Antimicrobial Activity. Biochemistry 58, 2447–2462

44. Anso, I., Basso, L. G. M., Wang, L., Marina, A., Paez-Perez, E. D., Jager, C., Gavotto, F., Tersa, M., Perrone, S., Contreras, F. X., Prandi, J., Gilleron, M., Linster, C. L., Corzana, F., Lowary, T. L., Trastoy, B., and Guerin, M. E. (2021) Molecular ruler mechanism and interfacial catalysis of the integral membrane acyltransferase PatA. Sci Adv 7, eabj4565

45. Scheithauer, L., Thiem, S., Schmelz, S., Dellmann, A., Bussow, K., Brouwer, R., Unal, C. M., Blankenfeldt, W., and Steinert, M. (2021) Zinc metalloprotease ProA of Legionella pneumophila increases alveolar septal thickness in human lung tissue explants by collagen IV degradation. Cell. Microbiol. 23, e13313

46. Jumper, J., Evans, R., Pritzel, A., Green, T., Figurnov, M., Ronneberger, O., Tunyasuvunakool, K., Bates, R., Žídek, A., Potapenko, A., Bridgland, A., Meyer, C., Kohl, S. A. A., Ballard, A. J., Cowie, A., Romera-Paredes, B., Nikolov, S., Jain, R., Adler, J., Back, T., Petersen, S., Reiman, D., Clancy, E., Zielinski, M., Steinegger, M., Pacholska, M., Berghammer, T., Bodenstein, S., Silver, D., Vinyals, O., Senior, A. W., Kavukcuoglu, K., Kohli, P., and Hassabis, D. (2021) Highly accurate protein structure prediction with AlphaFold. Nature 596, 583–589

47. Khan, F. I., Lan, D., Durrani, R., Huan, W., Zhao, Z., and Wang, Y. (2017) The Lid Domain in Lipases: Structural and Functional Determinant of Enzymatic Properties. Frontiers in bioengineering and biotechnology 5, 16–16

48. Gordon, V. M., and Leppla, S. H. (1994) Proteolytic activation of bacterial toxins: role of bacterial and host cell proteases. Infect. Immun. 62, 333–340

49. Kurmanova, A., Llorente, A., Polesskaya, A., Garred, Ø., Olsnes, S., Kozlov, J., and Sandvig, K. (2007) Structural requirements for furin-induced cleavage and activation of Shiga toxin. Biochem. Biophys. Res. Commun. 357, 144–149

50. Ogata, M., Fryling, C., Pastan, I., and FitzGerald, D. (1992) Cell-mediated cleavage of Pseudomonas exotoxin between Arg279 and Gly280 generates the enzymatically active fragment which translocates to the cytosol. J. Biol. Chem. 267, 25396–25401

51. van Deurs, B., and Sandvig, K. (1995) Furin-induced cleavage and activation of Shiga toxin. J. Biol. Chem. 270, 10817–10821

52. Veldhuizen, R., Nag, K., Orgeig, S., and Possmayer, F. (1998) The role of lipids in pulmonary surfactant. Biochim. Biophys. Acta 1408, 90–108

53. Laguna, R. K., Creasey, E. A., Li, Z., Valtz, N., and Isberg, R. R. (2006) A Legionella pneumophila-translocated substrate that is required for growth within macrophages and protection from host cell death. Proceedings of the National Academy of Sciences 103, 18745–18750

54. Ruiz-Albert, J., Yu, X. J., Beuzón, C. R., Blakey, A. N., Galyov, E. E., and Holden, D. W. (2002) Complementary activities of SseJ and SifA regulate dynamics of the Salmonella typhimurium vacuolar membrane. Mol. Microbiol. 44, 645–661

55. Roy, C. R. (2012) Vacuolar pathogens value membrane integrity. Proc. Natl. Acad. Sci. U. S. A. 109, 3197–3198

56. Miller, M. B., and Bassler, B. L. (2001) Quorum sensing in bacteria. Annu. Rev. Microbiol. 55, 165–199

57. Edelstein, P. H. (1981) Improved semiselective medium for isolation of Legionella pneumophila from contaminated clinical and environmental specimens. J. Clin. Microbiol. 14, 298–303

58. Bertani, G. (1951) Studies on lysogenesis. I. The mode of phage liberation by lysogenic Escherichia coli. J. Bacteriol. 62, 293–300

59. Bingel-Erlenmeyer, R., Olieric, V., Grimshaw, J. P. A., Gabadinho, J., Wang, X., Ebner, S. G., Isenegger, A., Schneider, R., Schneider, J., Glettig, W., Pradervand, C., Panepucci, E. H., Tomizaki, T., Wang, M., and Schulze-Briese, C. (2011) SLS Crystallization Platform at Beamline X06DA—A Fully Automated Pipeline Enabling in Situ X-ray Diffraction Screening. Crystal Growth & Design 11, 916–923

60. Vonrhein, C., Flensburg, C., Keller, P., Sharff, A., Smart, O., Paciorek, W., Womack, T., and Bricogne, G. (2011) Data processing and analysis with the autoPROC toolbox. Acta Crystallogr. D Biol. Crystallogr. 67, 293–302

61. Pape, T., and Schneider, T. R. (2004) HKL2MAP: a graphical user interface for macromolecular phasing with SHELX programs. Journal of Applied Crystallography 37, 843–844

62. Terwilliger, T. C., Grosse-Kunstleve, R. W., Afonine, P. V., Moriarty, N. W., Zwart, P. H., Hung, L.-W., Read, R. J., and Adams, P. D. (2008) Iterative model building, structure refinement and density modification with the PHENIX AutoBuild wizard. Acta Crystallographica Section D 64, 61–69

63. Emsley, P., Lohkamp, B., Scott, W. G., and Cowtan, K. (2010) Features and development of Coot. Acta Crystallogr. D Biol. Crystallogr. 66, 486–501

64. Winn, M. D., Ballard, C. C., Cowtan, K. D., Dodson, E. J., Emsley, P., Evans, P. R., Keegan, R. M., Krissinel, E. B., Leslie, A. G., McCoy, A., McNicholas, S. J., Murshudov, G. N., Pannu, N. S., Potterton, E. A., Powell, H. R., Read, R. J., Vagin, A., and Wilson, K. S. (2011) Overview of the CCP4 suite and current developments. Acta Crystallogr. D Biol. Crystallogr. 67, 235–242

65. Joosten, R. P., Long, F., Murshudov, G. N., and Perrakis, A. (2014) The PDB_REDO server for macromolecular structure model optimization. IUCrJ 1, 213–220

66. Afonine, P. V., Grosse-Kunstleve, R. W., Echols, N., Headd, J. J., Moriarty, N. W., Mustyakimov, M., Terwilliger, T. C., Urzhumtsev, A., Zwart, P. H., and Adams, P. D. (2012) Towards automated crystallographic structure refinement with phenix.refine. Acta Crystallogr. D Biol. Crystallogr. 68, 352–367

67. Burkhardt, A. (2016) Status of the crystallography beamlines at PETRA III. Eur. Phys. J. Plus 131

68. Mirdita, M., Schutze, K., Moriwaki, Y., Heo, L., Ovchinnikov, S., and Steinegger, M. (2022) ColabFold: making protein folding accessible to all. Nat Methods 19, 679–682

69. Flieger, A., Gongab, S., Faigle, M., Mayer, H. A., Kehrer, U., Mussotter, J., Bartmann, P., and Neumeister, B. (2000) Phospholipase A secreted by Legionella pneumophila destroys alveolar surfactant phospholipids. FEMS Microbiol. Lett. 188, 129–133

70. Schromm, A. B., Brandenburg, K., Rietschel, E. T., Flad, H. D., Carroll, S. F., and Seydel, U. (1996) Lipopolysaccharide-binding protein mediates CD14-independent intercalation of lipopolysaccharide into phospholipid membranes. FEBS Lett. 399, 267–271

71. Schrödinger, LLC. (2020) The PyMOL Molecular Graphics System, Version 2.4.

72. Jepras, R., Fitzgeorge, R., and Baskerville, A. (1985) A comparison of virulence of two strains of Legionella pneumophila based on experimental aerosol infection of guinea-pigs. Epidemiol. Infect. 95, 29–38

