## supplement for "Structure-function relationships underpin disulfide loop cleavage-dependent activation of *Legionella pneumophila* lysophosholipase A PlaA"

Figure S1: Details of *Legionella pneumophila* PlaA crystal structure determination.

Figure S2: Cross-eyed stereoplot of the substrate binding site.

Figure S3: Comparison of the PlaA crystal structure with an AlphaFold model.

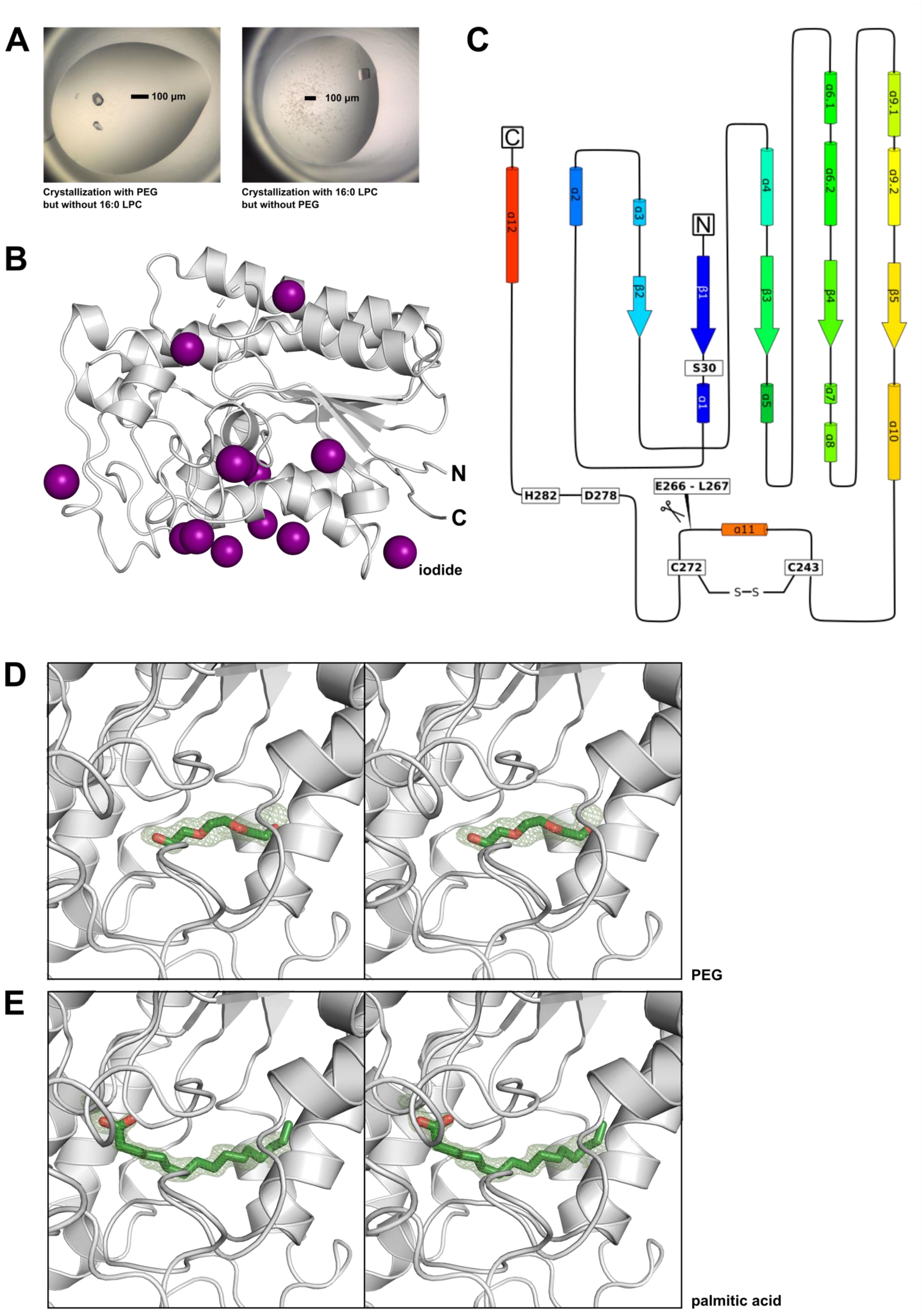


**Figure S1 Details of *Legionella pneumophila* PlaA crystal structure determination.** (*A*) PlaA crystals grown in 14.7% (w/v) PEG3350, 1% (w/v) PEG 1K, 0.5% (w/v) PEG200, 8.9% (v/v) tacsimate pH 5 and 200 mM $NH_4I$, 1 mM $CaCl_2$ and 1 mM $MgCl_2$ (left) or in 2.2 M $(NH_4)_2SO_4$ and 0.2 M $NH_4$-acetate after incubating PlaA at 7 mg/ml with 500 µM 16:0 LPC (right). (*B*) position of iodide anions (violet spheres) in PlaA (grey) after crystallization in 14.7% (w/v) PEG3350, 1%

(w/v) PEG 1K, 0.5% (w/v) PEG200, 8.9% (v/v) tacsimate pH 5, 200 mM $NH_4I$, 1 mM $CaCl_2$ and 1 mM $MgCl_2$ followed by cryoprotection in 10% (v/v) (2R,3R)-(−)-2,3-butandiole, 16% (w/v) PEG3350, 6% (v/v) tacsimate pH 5 and 920 mM $NH_4I$. Of 13 iodide anions contained in the final model (PDB entry 8A24), 11 produced anomalous density > 5 σ. (*C*) topology diagram of PlaA, colored from blue (N-terminus) to red (C-terminus). Colors correspond to Fig. 1A. (*D*) cross-eyed stereoplot of $|F_O\text{-}F_C|$ difference electron density of a PEG fragment bound to the substrate binding site of PlaA (PDB entry 8A25). The difference electron density is displayed at a level of 2.5 σ before incorporating PEG into the model. (*E*) cross-eyed stereoplot of $|F_O\text{-}F_C|$ difference electron density of palmitic acid bound to the substrate binding site of PlaA (PDB entry 8A26). The difference electron density is displayed at a level of 2.5 σ before incorporating palmitic acid into the model.

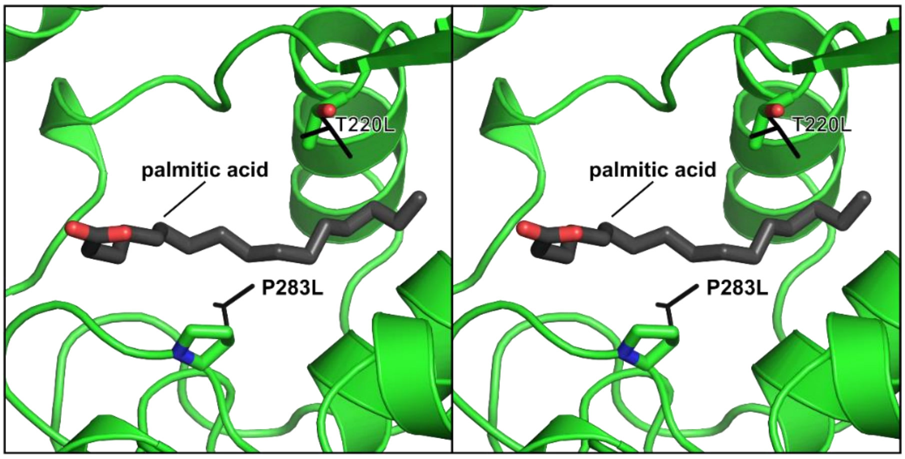


**Figure S2 Cross-eyed stereoplot of the substrate binding site.** Palmitic acid is shown in black. Mutation of P283 to leucine changed the substrate preference of PlaA to phospholipids containing shorter fatty acids, but not mutation of T220 to leucine."

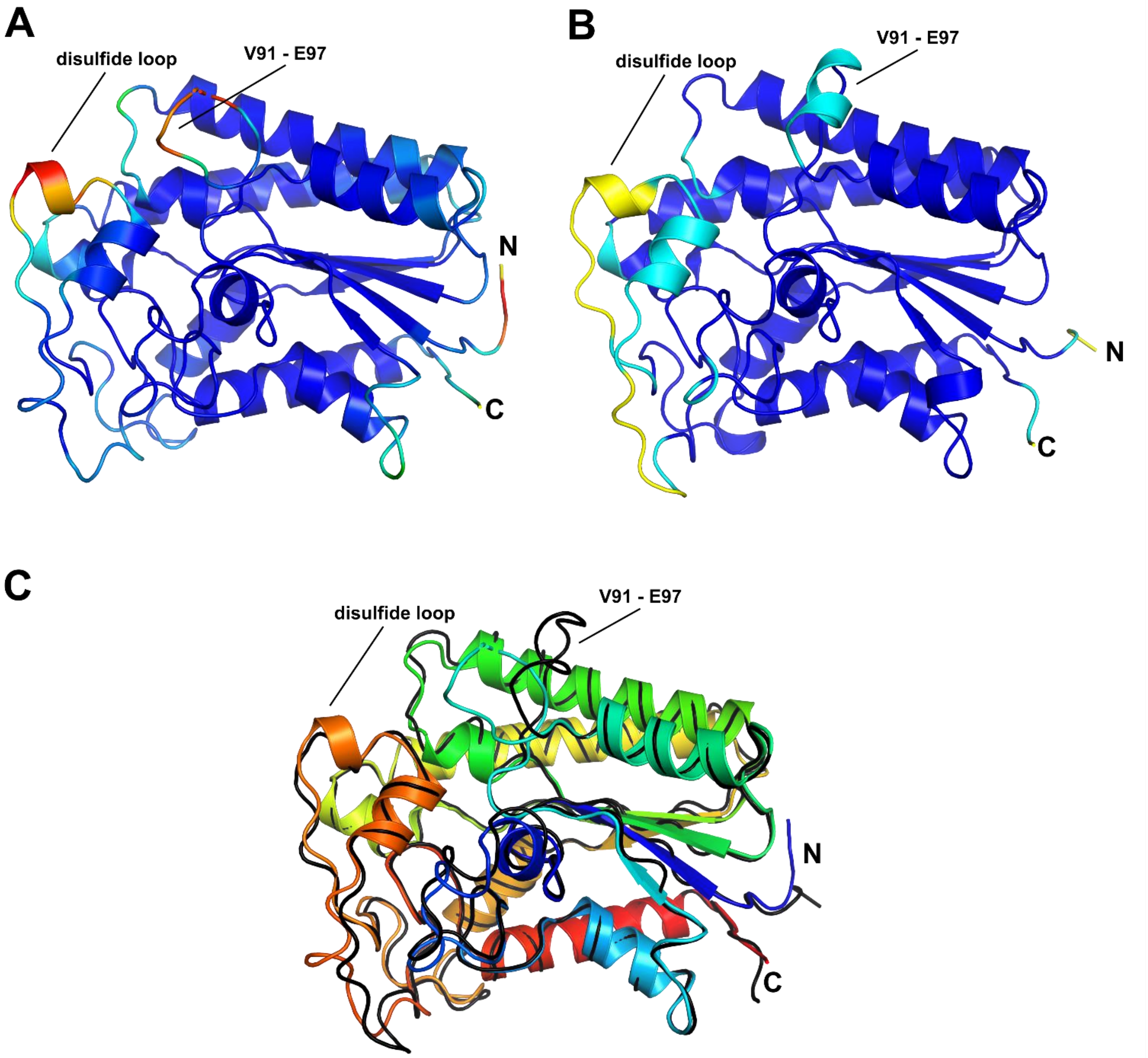


**Figure S3 Comparison of the PlaA crystal structure with an AlphaFold model.** (*A*) crystal structure of PlaA (complex with palmitate, PDB entry 8A26), colored according to B-factors (blue: low, red: high B-factor). (*B*) AlphaFold model of PlaA, colored according to pLDDT value (predicted local-difference distance test), a confidence measure for the correctness of structure prediction (46) (blue: pLDDT > 90 (high confidence), cyan: pLDDT > 70, yellow: pLDDT > 50, orange: pLDDT < 50 (low confidence)). Note that pLDDT values correlate to B-factors of the crystal structure. (*C*) superimposition of crystal structure and AlphaFold model (superimposition performed with PyMol (71)). The RMSD value is 0.77 Å$^2$ for 279 (out of 289 possible) residues.
